# Modeling and dissecting bidirectional feedback in gene-metabolite systems using the CausalFlux method

**DOI:** 10.64898/2026.04.10.717623

**Authors:** Nilesh Subramanian, S Pavan Kumar, Raghunathan Rengaswamy, Nirav Pravinbhai Bhatt, Manikandan Narayanan

## Abstract

Predicting cellular behaviors, a central task in systems biology and metabolic engineering, can be enhanced through integrative modeling of processes such as gene regulation and metabolism. Information flow from gene regulation (modeled via a gene regulatory network) to metabolism (modeled via a genome-scale metabolic model) is well-studied, but the reciprocal regulation of genes by metabolites is less explored. We introduce CausalFlux, a method that models bidirectional feedback between genes and metabolites, in order to predict steady-state reaction fluxes under wild-type (WT) or perturbed (e.g., gene knockout/KO) conditions. CausalFlux does so by iteratively performing causal surgery on a Bayesian gene regulatory network and constraint-based analysis of a coupled metabolic model. CausalFlux enabled us to assess the impact of two-way feedback in several testbed models and real-world biological systems by comparing its predictions to those of TRIMER, a state-of-the-art model of gene-to-metabolite one-way feedback. Incorporating bidirectional feedback, as in CausalFlux, improved the Spearman correlation between actual and predicted fluxes in 92% of the 39 distinct simulation conditions relative to TRIMER. For predicting growth/no-growth phenotype following single-gene KOs in *E. coli*, CausalFlux achieved a balanced accuracy of 0.79 in identifying essential genes, and TRIMER achieved 0.71 for the same task, again highlighting the importance of modeling two-way feedback. In ablation studies that further dissect the role of specific metabolite→gene feedback edges in *E. coli*, the F1 scores of gene essentiality predictions decreased by 7.5% and 13% upon ablation of feedback edges from any metabolite to the *crp* gene and the 10 metabolic feedback genes with the highest influence on the KO genes, respectively. Finally, we highlight the application of CausalFlux to predict the essentiality of several hundred genes under different media conditions. Overall, our findings show that CausalFlux can crucially utilize information on feedback metabolites to predict trends in reaction fluxes and qualitative (growth/no-growth) outcomes; thereby encouraging future systems modeling efforts to carefully incorporate not only gene-to-metabolite but also metabolite-to-gene interactions.

**Availability:** Code pertaining to the CausalFlux method, and its benchmarking and application is publicly available at: https://github.com/BIRDSgroup/CausalFlux.

**Author summary:** The myriad processes within a living cell, such as gene regulation or metabolism, are tightly interconnected. Modeling these interconnected processes can offer a deeper mechanistic understanding of cellular behaviors, as well as guide efforts that engineer the metabolic output of a cell. In this work, we develop a novel integrated model of gene regulation and metabolism that incorporates bidirectional feedback between these two processes, via the concept of metabolite-induced causal surgery on a gene regulatory network and gene-induced constraints on the fluxes of metabolic reactions. Our model, which we call CausalFlux, represents an advance over most existing models that capture just the one-way gene-to-metabolism feedback (i.e., genes coding for enzymes that control metabolic reactions). Our CausalFlux methodology opens up an unique opportunity to quantify the impact of two-way feedback in gene-metabolite systems, via comparison of CausalFlux’s predictions to those of TRIMER, a published model incorporating one-way feedback alone. For predicting reaction fluxes in testbed models and essential genes in *E. coli*, quantitative comparison of the performance of CausalFlux vs. TRIMER showed that accounting for two-way feedback leads to more accurate and biologically meaningful predictions. CausalFlux also enabled us to quantify the effect of two-way feedback by comparing prediction performance before and after ablation of certain feedback edges from metabolites to genes. Overall, our findings highlight the importance of modeling gene regulation and metabolism as two-way interconnected systems within a living cell, and encourage future works to incorporate gene↔metabolite feedback into their analyses.

## Introduction

Thousands of genome-scale metabolic models (GSMM) have been constructed in the past two decades. These network-based models comprehensively represent an organism’s metabolism, capturing the network of biochemical reactions as a stoichiometric matrix and gene-protein-reaction (GPR) rules that govern the activity of specific metabolic reaction. Very often, the gene regulatory state of the cell inferred via transcriptomic (gene expression) measurements and/or knowledge of the gene regulatory network (GRN) is integrated as a constraint to improve GSMMs’ predictions. Existing methods that integrate gene regulatory constraints into GSMMs can be broadly classified into three categories [1]: (i) Integration through transcriptomics data: These methods filter out (or capture) certain metabolic reactions by evaluating the GPR rules using the gene expression evidence, thereby representing the metabolic reactions as either ON (active) or OFF (inactive). Examples include context-specific model-building algorithms such as GIMME [2], and Fastcore [3]. One of the major limitations of these approaches is their inability to predict fluxes when gene expression data corresponding to the KO (Knockout) perturbation of interest is not available; (ii) Integration through Boolean GRNs: Methods like rFBA [4, 5] and SR-FBA [6, 7] represent gene expression states as ON/OFF and model regulatory programs using a network of Boolean relations involving these genes. They also model metabolic regulation of genes into this Boolean network formulation to capture bidirectional gene-metabolite feedback, but they require extensive literature searches to define the Boolean relations/rules and can be challenging to scale up to a genome-wide model of complex systems; and (iii) Probabilistic representation of the GRN: Methods like PROM [8] and TRIMER [9] infer the conditional probabilities of genes (e.g., target genes) conditioned on the state of their regulating genes (e.g., corresponding transcription factors). The advantage of TRIMER over PROM is the use of a GRN reconstructed from expression data and prior-known gene regulatory interactions. However, these methods do not account for the feedback from the metabolic network to the GRN.

Multi-omics studies on *E. coli* and yeast have revealed the presence of certain metabolites that regulate the expression of genes in the GRN [10], thereby exerting feedback on gene expression and creating crosstalk between metabolic and regulatory systems. Carthew provides examples of metabolites acting as signaling molecules to regulate genes in the GRN [11]. Kumar et al. utilize this knowledge of metabolite feedback to construct different network modules of the GRN, both with and without feedback, to understand the underlying causal structure of the GRN in *E. coli* and *B. subtilis* [12]. These studies emphasize that incorporating metabolic feedback into the GRN is crucial to obtain a tightly integrated model of the cellular subsystems involved, and thereby encourages extending current efforts that mostly model one-way feedback from gene regulation to metabolism to explicitly capture bidirectional feedback.

Towards addressing the above research gap, this study makes the following broad contributions:

i. Methodological: We propose ‘CausalFlux’, an integrated framework that captures the complex interactions within and between the metabolic and gene networks (see Fig. 1[A–C]). CausalFlux specifically models the bidirectional feedback between gene regulation and metabolism in an iterative fashion. CausalFlux’s unique contribution is to model metabolite-to-gene interactions, i.e., feedback metabolites that regulate gene activity, via the techniques of iterative causal surgery on a Bayesian network representation of GRN, and constraint-based analysis of a coupled metabolic model (see Fig. 1[B–C]).
ii. Conceptual: Equipped with CausalFlux, we systematically quantified the impact of bidirectional feedback in a cellular system by comparing CausalFlux’s predictive performance to that of TRIMER, and via ablation of certain feedback edges. We performed the comparative analyses on testbed models (TMs) with simulated steady-state fluxes, and a real-world *E. coli* dataset with ground-truth information on growth/no-growth phenotype of 798 single-gene knockouts (KOs; see Fig. 1[D–E]); and ablation studies on the *E. coli* dataset.
iii. Application: We highlight an application of CausalFlux to predict gene essentiality of several hundred genes under different growth media conditions (minimal to rich media). Such prospective predictions can aid metabolic engineering efforts by prioritizing single-gene KOs that yield the desired phenotype of interest.

**Fig 1.**
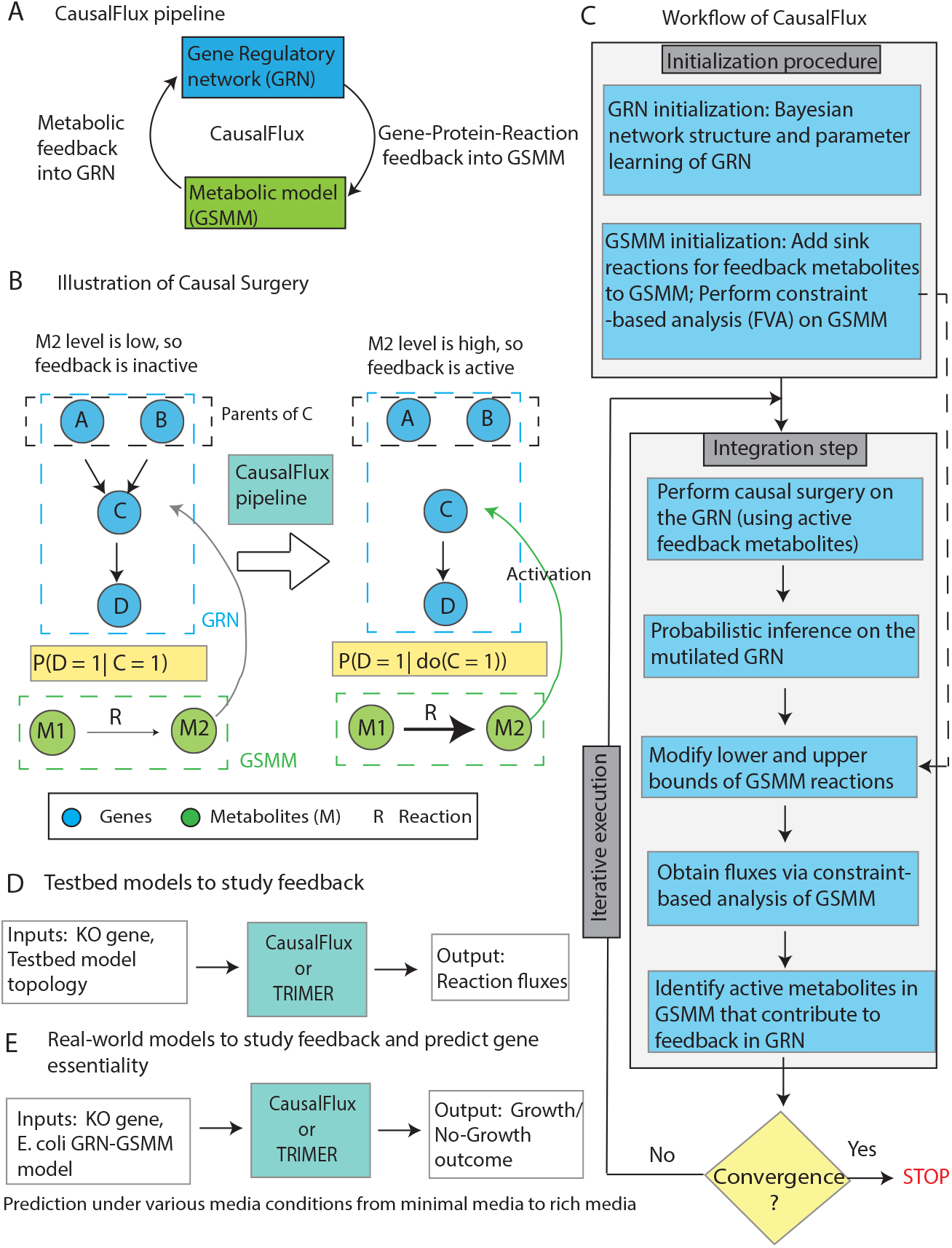
Overview of our CausalFlux method: (A) CausalFlux integrates GRN model and GSMM using prior knowledge on two-way feedback interactions between them, in an iterative fashion. (B) A key step in CausalFlux is performing causal surgery on a conditioning variable like gene “C” (which may be a transcription factor in the GRN) – incoming edges from the parents of “C” (“A” and “B”) are severed to obtain a mutilated GRN, and “C”‘s value forced to 1 to reflect activation feedback from M2 metabolite. Conditional probability of “D” before and after causal surgery is shown. (C) CausalFlux flowchart (see Algorithm 1 and Methods for details; FVA is flux variability analysis). (D & E) Comparison of predictive performance of CausalFlux vs. one-way feedback method TRIMER (or GIMME) on: (D) three testbed GRN+GSMM integrated models, with fluxes simulated using ODEs under various exchange rates and gene KOs (ODE is Ordinary Differential Equation); and (E) real-world *E. coli* model, with growth/no-growth outcomes measured under 798 single-gene KOs.

Overall, the results from the analyses and application above showed that the CausalFlux method that we developed can crucially utilize information on feedback metabolites to predict trends in reaction fluxes and qualitative (growth/no-growth) outcomes; thereby encouraging future systems modeling efforts to carefully incorporate not only gene-to-metabolite but also metabolite-to-gene interactions.

## Results

As seen in Fig. 1, CausalFlux integrates GRN and GSMM iteratively, through bidirectional feedback to perform causal surgery on the GRN, followed by constraint-based analysis of the GSMM. After convergence, CausalFlux predicts the steady-state reaction fluxes in the GSMM by implementing Flux Balance Analysis (FBA) and Flux Variability Analysis (FVA), yielding FBA solutions as well as minimum and maximum FVA fluxes (FVA min and FVA max) for each reaction. We evaluate CausalFlux using multiple testbed models (TMs) (see Fig. 2A for the topology of TM 1; topologies of TM2 and TM3 are in S1 Fig), and real-world datasets, as described in the subsequent sections.

**Fig 2.**
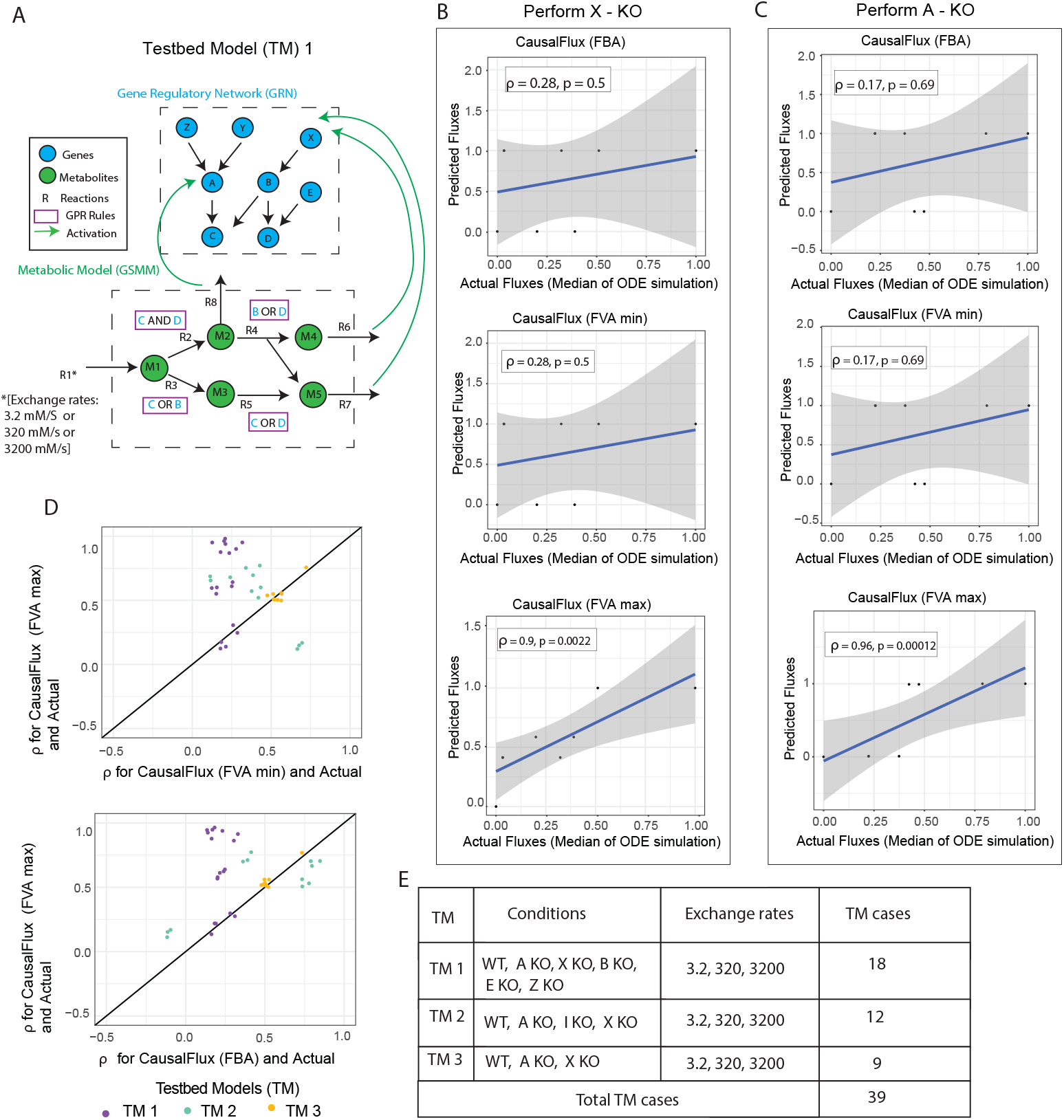
FVA max as best CausalFlux variant on TMs: (A) Topology of TM1 (integrated GRN-GSMM system; see S1 Fig. for the topology of TM2 and TM3). (B-C) Scatter plots between the normalized predicted and actual fluxes for TM1 under the 320 exchange rate for “X” and “A” gene KOs, where predictions are by CausalFlux’s FBA and FVA (min and max) variants. Spearman’s *ρ* and associated p-value are also shown atop each plot. (D) Scatter plots (with jitter of 0.05 units) of the *ρ* computed between different CausalFlux variants’ predicted and actual fluxes for each of the 39 TM cases. (E) Different simulation scenarios/cases for each TM and associated correlation tests performed (points in plot D) are listed.

### Evaluating CausalFlux on testbed models

To evaluate our method, we computed the Spearman correlation (*ρ*) between ODE-simulated ground-truth fluxes (see Methods for flux simulations) and predicted fluxes across 39 TM cases (see Fig. 2E). For CausalFlux, we selected the best-performing variant among FBA, FVA min, and FVA max. We then compared these results with predictions from other methods applied to TMs, namely GIMME (FVA max) and TRIMER (sFBA). GIMME (FVA max) was chosen based on its superior performance relative to GIMME (FVA min) and GIMME (FBA), while sFBA was the best-performing TRIMER variant as reported in the original study [9]. An overview of the simulation-based benchmarking pipeline is shown in S2B Fig.

### FVA max achieves superior performance among CausalFlux variants

To identify the best-performing CausalFlux variant among FBA, FVA max, and FVA min, we compared predicted reaction fluxes with ground-truth ODE-simulated fluxes. The median of the ODE flux simulations was taken as the ground truth (see Methods for details). Scatter plots comparing actual and predicted fluxes for FBA, FVA min, and FVA max variants of CausalFlux for “X” and “A” gene knockouts in TM1 are shown in Fig. 2B and C. These plots indicate that the FVA max variant performs better than FVA min and FBA, as reflected by higher correlation (*ρ*) values for both knockout cases. We next expanded our evaluation from these two KO cases to all 39 TM cases. In a majority of these TM cases, we found that the predicted fluxes of CausalFlux (FVA max) showed higher *ρ* values (i.e., higher correlation with actual fluxes) than other CausalFlux variants, FVA min and FBA (Fig. 2D). Specifically, FVA max outperformed FVA min and FBA in 92% and 84% of cases, respectively. Together, these results indicate that the FVA max variant of CausalFlux consistently outperforms its FVA min and FBA counterparts, and was therefore selected for subsequent comparative evaluations. Refer to Supplementary Data 1 to view the (*ρ*) computed for the various methods for all the WT/KO cases done for all the TMS under various exchange rates.

### CausalFlux (FVA max) is better than or comparable to other state-of-the-art or baseline methods

To compare the performance of CausalFlux predictions with predictions from other state-of-the-art baseline methods (which do not explicitly model two-way feedback as mentioned earlier), we looked at the following aspects: (I) Overall CausalFlux performance in predicting fluxes of all reactions across all 39 TM cases; (II) CausalFlux performance in predicting the flux of the biomass reaction across the 39 TM cases; and (III) Dissecting the role of feedback in the performance improvements.

#### Overall performance of CausalFlux across all the reactions in TMs

The correlation (*ρ*) computed between actual and predicted fluxes for GIMME (FVA max), GIMME (FVA min), and GIMME (FBA) were largely comparable, with FVA max performing slightly better; therefore, GIMME (FVA max) was selected for subsequent comparisons (see S3 Fig). Using this configuration, CausalFlux (FVA max) outperformed both TRIMER and GIMME (FVA max), achieving higher *ρ* values between predicted and actual fluxes in 92% and 69% of the total TM cases, respectively (see Fig. 3[A–B]). This performance trend persists across varying exchange rates (see Fig. 3[C–D]), suggesting that CausalFlux’s superior performance is robust against changes in input exchange rates.

**Fig 3.**
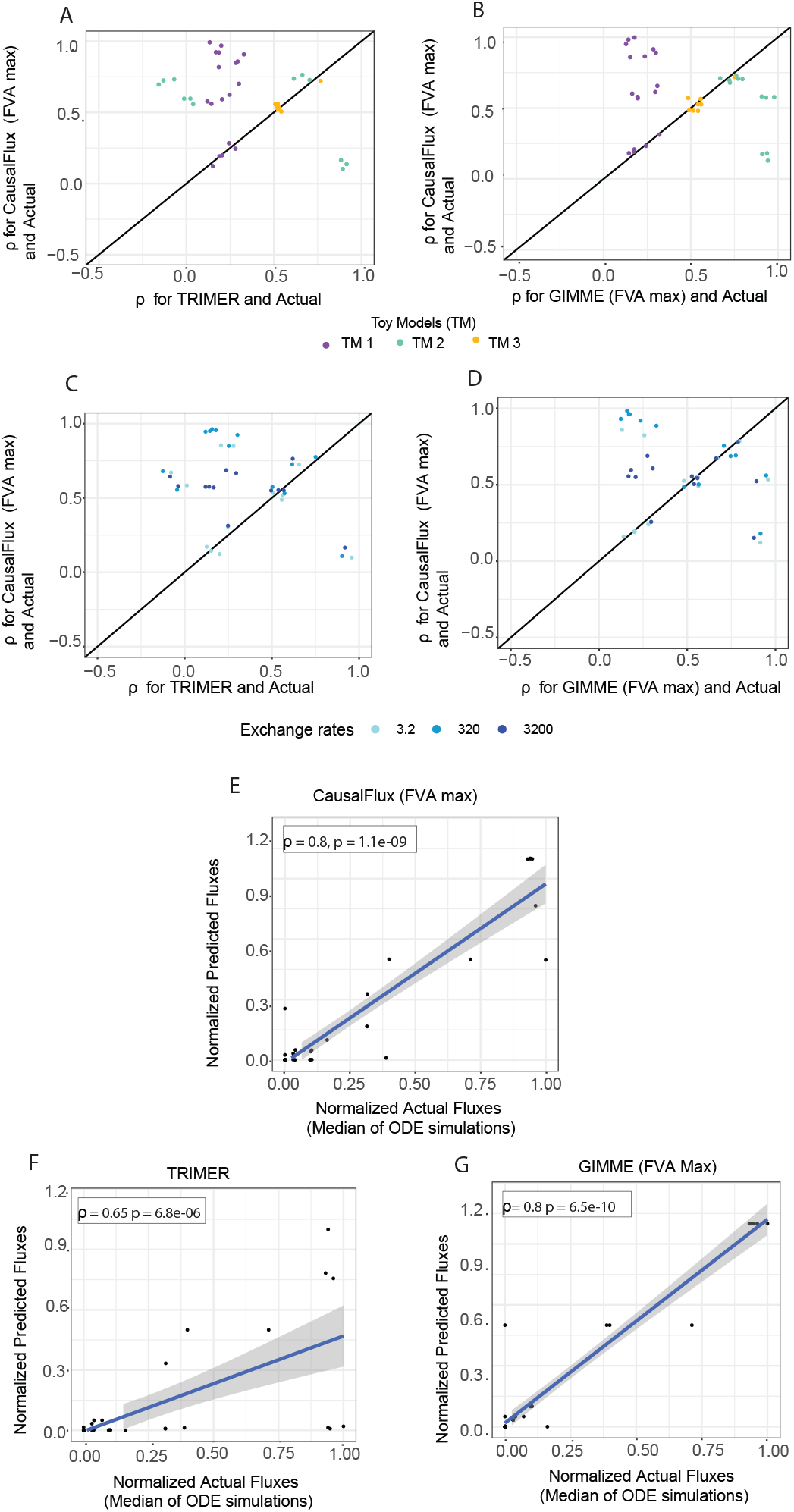
Comparing CausalFlux, TRIMER and GIMME on TMs: (A) Scatter plot (with jitter of 0.05 units) of *ρ* computed between CausalFlux (FVA max) and actual (vs.) TRIMER and actual for all the WT/KO cases across all TMs under different exchange rates. (B) Similar plot as (A) but with TRIMER replaced by GIMME (FVA max). (C) and (D) are the same as (A) and (B), respectively, but the points are colored based on the exchange rates. (E), (F), and (G) Actual vs predicted (CausalFlux (FVA max), TRIMER, and GIMME (FVA max) respectively) fluxes of the biomass reaction across the WT/KO cases in all TMs under different exchange rates.

#### Performance of CausalFlux for the biomass reaction in TMs

The *ρ* between biomass fluxes predicted by CausalFlux (FVA max) and the corresponding actual fluxes is 0.80, which is higher than the *ρ* of 0.65 obtained using TRIMER predictions (see Fig. 3[E–G]). This indicates that CausalFlux (FVA max) better captures the trends in biomass fluxes across TM wild-type and knockout cases and across all exchange rates. CausalFlux (FVA max) and GIMME (FVA max) exhibit comparable performance, with GIMME showing a slight advantage in terms of p-value. This difference may be attributed to the additional information available to GIMME in the form of gene expression data specific to the considered gene knockout. In contrast, CausalFlux achieves comparable performance without relying on such KO-specific expression data.

#### Effect of feedback-imparting iterations on the CausalFlux outcome

To better understand the role of feedback in our method, we examined whether the actual fluxes were more strongly correlated with the fluxes predicted from successive iterations of CausalFlux (compared to its earlier iterations where feedback is incorporated to a lesser extent). *ρ* was computed between predictions from iteration 1 and the actual fluxes across all 39 TM cases, and similarly between predictions from iteration 2 and the actual fluxes. We observed that in certain cases, such as the “WT” and “X” knockouts in TM1 and TM2, respectively, feedback improved performance, as reflected by increased *ρ* values in successive iterations (see S4 Fig). In contrast, for most of the other cases, the data points lie close to the *x* = *y* line, indicating that feedback had little effect. In a few specific scenarios, such as the “I” knockout in TM2 and the 3200 exchange rate condition in TM1, feedback appeared to have a negative impact.

Together, all of the above observations suggest that the influence of feedback is context-dependent, and a more careful examination in real-world settings is required to understand its effects fully, which we perform next.

### Real-world evaluation: Using CausalFlux to predict growth/no-growth phenotype in *E. coli*

We applied CausalFlux and TRIMER to computationally model 798 single-gene KOs in *E. coli* (see Methods on implementation) to predict fluxes. For each gene KO, if the predicted flux for the biomass reaction is 0, then a “no-growth” phenotype is predicted (i.e., the gene is predicted as an essential gene). If the gene KO leads to a prediction of non-zero biomass flux, then we predict a growth phenotype (i.e., that the gene is non-essential). Of all single-gene knockouts considered, 47 genes showed zero growth upon knockout in the Keio collection and were therefore treated as ground-truth essential; the rest were called ground-truth non-essential genes [13].

CausalFlux performs better in predicting the no-growth cases (essential genes) compared to TRIMER, and a random classifier, as seen in Fig. 4A. Here, the random classifier refers to a classifier that predicts all the instances as the positive class (essential or no-growth cases). CausalFlux predictions were better than those of TRIMER in terms of F1 score and balanced accuracy (see Fig. 4A). Of the 47 ground-truth essential genes, 29 were accurately predicted by CausalFlux. While TRIMER accurately predicted 22 out of the 47 essential genes, it also predicted a larger number of false positives.

**Fig 4.**
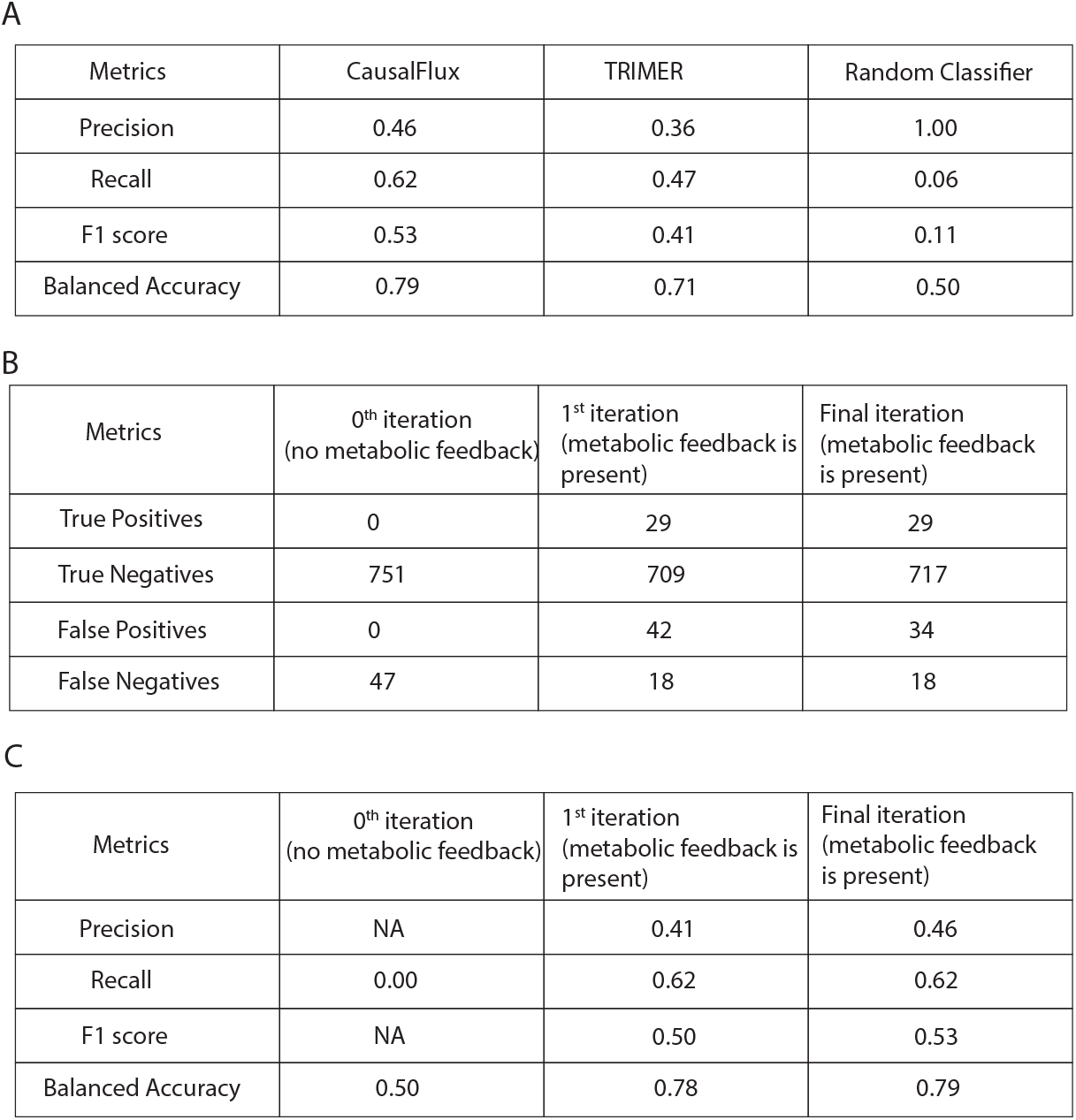
Comparing CausalFlux and TRIMER on *E. coli* datasets: (A) Comparing the performances of CausalFlux, TRIMER, and a random classifier for predicting growth/no growth in 798 single-gene KO cases in *E. coli*. (B) Comparing the predictions and (C) metrics for CausalFlux across 0^*th*^, 1^*st*^ iterations and after the algorithm had converged to show the effect of metabolic feedback. Note: True positives: essential genes predicted as essential; True negatives: non-essential genes predicted as non-essential; False positives: non-essential genes predicted as essential; and False negatives: essential genes predicted as non-essential.

To assess the impact of metabolic feedback on CausalFlux predictions, we compared the results and corresponding metrics at the end of three iterations of the algorithm: the 0^*th*^ iteration (without any metabolic feedback), the 1^*st*^ iteration (after one round of feedback incorporation), and the final iteration (the final converged state after multiple feedback iterations), as shown in Fig. 4[B–C]. An improvement in both precision and F1 score can be clearly observed across these iterations, indicating that incorporating metabolic feedback enhances the performance of CausalFlux. We could also observe changes in true negatives (from 751→709→717) and false positives (from 0→42→34) over iterations. These findings show that feedback-imparting iterations are useful to enhance the performance of CausalFlux in real-world *E. coli* benchmarks.

### Ablation of metabolic feedback edges to genes highlights the role of feedback in regulating growth

To better understand the influence of metabolites on gene regulation, we performed a series of ablation studies in which subsets of feedback edges from metabolites to genes were removed (Fig. 5A). We focused on ablation study of metabolic feedback gene(s) with high influence on single-gene KOs. Influence of a gene A is quantified by the number of 798 single-KO genes that are in the 0-hop or 1-hop neighbourhood of A in the GRN (see Table S4 for a list of metabolic feedback genes sorted by their influence). The different ablation studies and the number of genes misclassified as essential after ablation is summarized in Fig. 5A. Notably, several of these genes were immediate (1-hop) neighbors of metabolic feedback genes.

**Fig 5.**
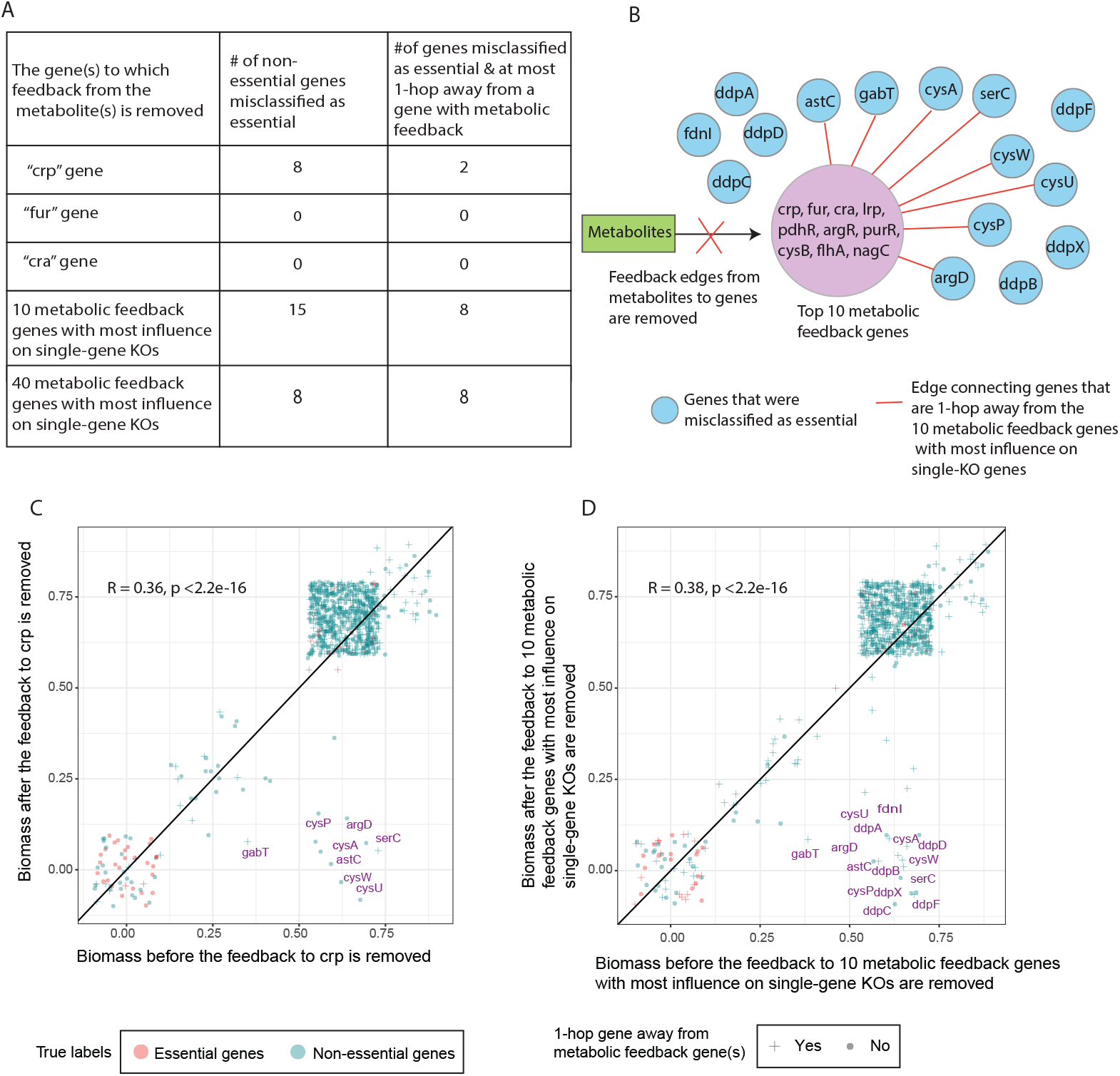
Exploring the role of metabolic feedback through ablation studies: (A) Five different ablation studies were performed by removing the metabolite feedback edges to gene(s) that have high influence on single-gene KOs. Information on genes misclassified as essential after ablation are also shown. (B) Illustration of the fourth ablation study (fourth row) shown in panel A. (C and D) Scatter plot (with default jitter) showing the biomass of the 798 single-gene KOs before vs. after removal of feedback edges to: the gene *crp* (panel C); and the 10 metabolic feedback genes with the most influence on single-gene KOs (panel D).

In the first ablation study, we focused on the metabolic feedback gene with the highest influence on the 798 single-KO genes, viz., gene *crp*. Note that *crp* is a global transcriptional regulator that regulates multiple genes in the catabolic and anabolic pathways in *E. coli* [14]. After ablation of the feedback edges to *crp*, the biomass remained the same as before ablation under single-KO of most of the 798 genes (as seen in Fig. 5C). However, interestingly, certain genes such as *astC, gabT, cysA, serC, cysW, cysP, cysU* and *argD* (labeled in purple in Fig. 5C) were misclassified as essential genes after ablation but not before; and out of these genes, *gabT* and *serC* are children of (1-hop from) *crp* in the GRN, suggesting that allowing metabolic control of *crp* properly modulates its effect on these two downstream genes. A deeper analysis of reaction fluxes under *gabT* KO, comparing the model incorporating metabolic feedback of *crp* with the ablated model without this feedback, revealed that 123 reactions exhibited non-zero flux in the former model and zero flux in the latter ablated model. Among these, 27 reactions showed differences in *GPR* probabilities between the two models, with 24 of these reactions directly regulated by genes downstream of *crp*. Notably, several of these reactions are part of key metabolic pathways that produce metabolites that directly contribute to the biomass reaction, and could explain why we see no growth in the ablated model (examples of such reactions include “Succinyldiaminopimelate transaminase”, “O-Phospho-4-hydroxy-L-threonine:2-oxoglutarate aminotransferase”, and “Nicotinate-nucleotide diphosphorylase”, which are involved in biosynthesis of lysine, serine, and de novo NAD respectively [15, 16, 17]; see Supplementary Data 2 for other reaction names). A similar trend was observed under *serC* KO. That is, 144 reactions exhibited zero flux in the ablated model, of which 29 reactions showed differences in *GPR* probabilities between the two models, and 24 of these 29 reactions were directly regulated by genes downstream of *crp*. Consistent with the overall findings above, previous studies have reported that *crp* regulates members of the *gab* gene family as well as *serC*, which are involved in amino acid biosynthesis pathways such as lysine and serine biosynthesis [18, 19].

On the other hand, repeating the same ablation analysis separately on *fur* and *cra*, the metabolic feedback genes with the 2^*nd*^ and 3^*rd*^ highest influence on single-KO genes, respectively, did not affect biomass predictions (see S5[A–B] Fig). These observations led us to perform ablation analysis on a set of 10 metabolic feedback genes with the most influence on single-gene KOs to check the overall effect (see Fig. 5B, and Methods). After this ablation, CausalFlux misclassified 15 genes (blue nodes in Fig. 5B and labeled points in Fig. 5D) as essential, out of which 8 genes were 1-hop away from (at least one of) the 10 metabolic feedback genes considered (see also fourth row in Fig. 5A). A similar observation was also made from a similar ablation analysis performed on 40 metabolic feedback genes with the most influence on single-gene KOs (see S5C Fig.). In this case, 8 non-essential genes were misclassified as essential, all of which were 1-hop neighbours of (at least one of) the 40 metabolic feedback genes. These ablation-induced misclassifications (including the proximity of genes misclassified as essential to highly influential metabolic feedback genes) indicates that the metabolic feedback of multiple genes has a significant overall effect on biomass.

As a final ablation analysis, we quantitatively assessed the contribution of metabolite-to-gene feedback to biomass by comparing CausalFlux’s performance metrics before vs. after ablation (see Table S5). Excepting the two ablations involving *fur* and *cra*, the percentage decrease in F1 score of the other three ablated models relative to the non-ablated model ranged from 7.5% to 13%. The ablation studies overall highlighted multiple scenarios where metabolite-to-gene feedback edges play a crucial role in proper modeling of cell growth.

### Benchmarking CausalFlux with rFBA and PROM datasets

We also benchmarked our method against two other older methods, rFBA [5] and PROM [8]. Chandrasekaran et al. in their work compare their work with rFBA by predicting the biomass of E.coli under 100 media conditions for 15 gene KOs and ground truth biomass values. The 100 media conditions comprise 58 carbon sources, 31 nitrogen sources, and 11 double-nitrogen sources. We used these results to compare the performances of CausalFlux run on rFBA and PROM datasets for the same scenario (See Methods for a more detailed description of the datasets). Consistent with our real-world analyses, CausalFlux classifies genes as essential or non-essential based on zero versus non-zero predicted biomass, and we maintain this definition here. In contrast, for rFBA and PROM, a gene is classified as non-essential if the predicted biomass is at least 5% of the wild-type (WT) biomass; otherwise, it is classified as essential.

The biomass predictions of CausalFlux across 100 media conditions were categorised as no growth and growth (in PROM and rFBA this is referred to as lethal/non-lethal cases, respectively). The scatter plot in Fig. 6A compares the accuracy of CausalFlux predictions with the ground truth (binary data of no-growth and growth outcomes) against the corresponding accuracy of rFBA predictions for each media condition across 15 gene knockouts. Each red dot in the figure represents one media condition. The plot shows that most points lie slightly below the diagonal, indicating that the accuracy of CausalFlux predictions tends to be marginally higher than that of rFBA. As summarised in Fig. 6C, CausalFlux outperforms rFBA in 56 out of 100 cases, performs equally in 17 cases (points on the diagonal), and has lower performance in 27 cases (points above the diagonal). The performance breakdown for each of the three major sources is also provided in Fig. 6C. In comparison with PROM, most of the points lie along the diagonal in Fig. 6B, indicating comparable performance (89 out of 100 cases, as shown in Fig. 6D). However, in 9 out of 100 cases, CausalFlux performs slightly lower than PROM.

**Fig 6.**
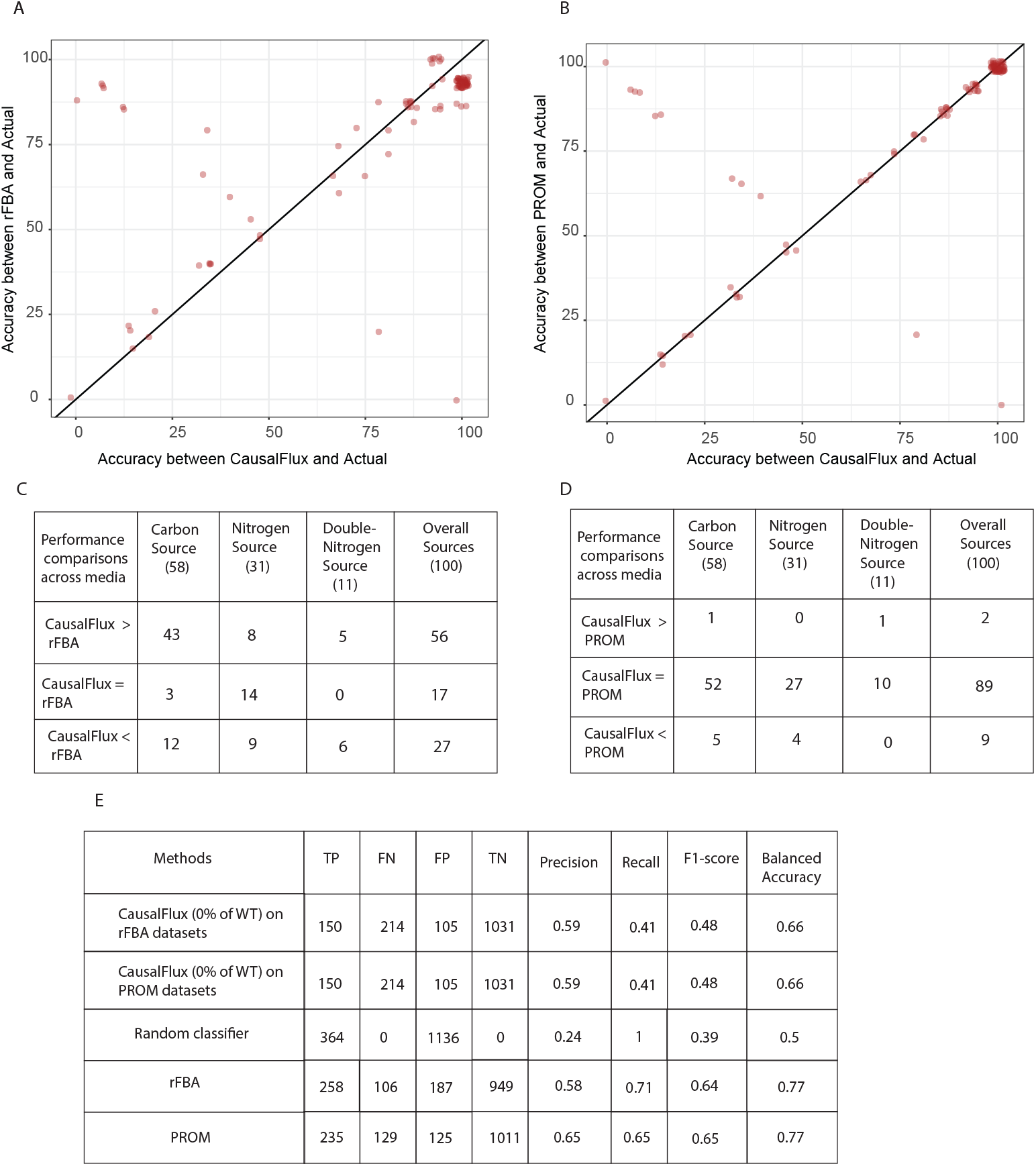
Benchmarking CausalFlux on PROM and rFBA datasets: (A) Scatter plot (with jitter 1.75 units) between the accuracy computed between CausalFlux predictions and Actual vs. the accuracy between rFBA predictions and Actual for the 100 media conditions. (B) Scatter plot (with jitter 1.75 units) between the accuracy computed between CausalFlux predictions and Actual vs. the accuracy between PROM predictions and Actual for the 100 media conditions. (C) Performance comparisons of CausalFlux and rFBA for the various Carbon sources, Nitrogen sources and Double-Nitrogen sources and overall 100 media conditions. (D) Performance comparisons of CausalFlux and PROM for the various Carbon sources, Nitrogen sources and Double-Nitrogen sources and overall 100 media conditions. (E) Comparison of classification metrics (TP: true positives [essential], FN: false negatives, FP: false positives, TN: true negatives [non-essential]) for CausalFlux applied to the rFBA and PROM datasets (usual definition of essential/non-essential), along with Random Classifier, rFBA, and PROM predictions across 1500 data points (100 media conditions *×* 15 gene KOs).

We also analyzed the individual 1,500 data points (100 media conditions × 15 gene KOs), and various performance metrics are reported for the predictions of CausalFlux when applied to the rFBA and PROM datasets individually, as well as for a random classifier that predicts all points as no-growth cases, and for the rFBA and PROM predictions themselves (see Fig. 6E). CausalFlux outperforms the random classifier in both cases. Compared to rFBA and PROM, the F1 scores of our approach are slightly lower, primarily due to reduced recall values. However, our predictions exhibit fewer false positives and a higher number of true negatives relative to the other methods, indicating better specificity (0.91 for both CasualFlux predictions, 0.84 for rFBA and 0.89 for PROM).

We also varied the definition of no-growth and growth cases (based on predicted biomass) using thresholds of 5% and 50% of the wild-type biomass values. Chandrasekaran et al. previously defined no-growth cases as those with biomass less than 5% of the wild-type biomass in their comparison with rFBA. Following this, we used both 5% and 50% thresholds to explore the effect of this criterion and generated the corresponding scatter plots and summary tables (Fig. 6) for CausalFlux predictions on the rFBA and PROM datasets, with additional results provided in S6 and S7 Fig. In the 5% scenario, the results of CausalFlux on rFBA were identical to those of the default zero/non-zero case described earlier. When applied to the PROM datasets, PROM outperformed CausalFlux in a greater number of conditions, and this trend persisted in the 50% scenario. Nevertheless, across all three growth definitions, CausalFlux performed better than rFBA in terms of the number of conditions where it achieved higher accuracy.

### Applying CausalFlux to predict growth/no-growth phenotype under different media conditions

We applied CausalFlux to perform 798 single-gene KOs in *E. coli* under other media conditions, namely M9 minimal medium and Tryptic Soy Broth (TSB). Knockout analysis showed that while growth (nonzero biomass) was often maintained in both LB and M9 media, M9 exhibited a higher number of zero-growth cases, indicating a greater number of essential gene knockouts (Fig. 7[A–B]). This observation is expected, as M9 is a minimal media. In comparison with LB and TSB predictions, the prediction under TSB conditions showed slightly more zero-growth cases than LB. Overall, M9 media resulted in the highest number of predicted essential gene knockouts (133), followed by TSB (93) and LB (63) (Fig. 7C).

**Fig 7.**
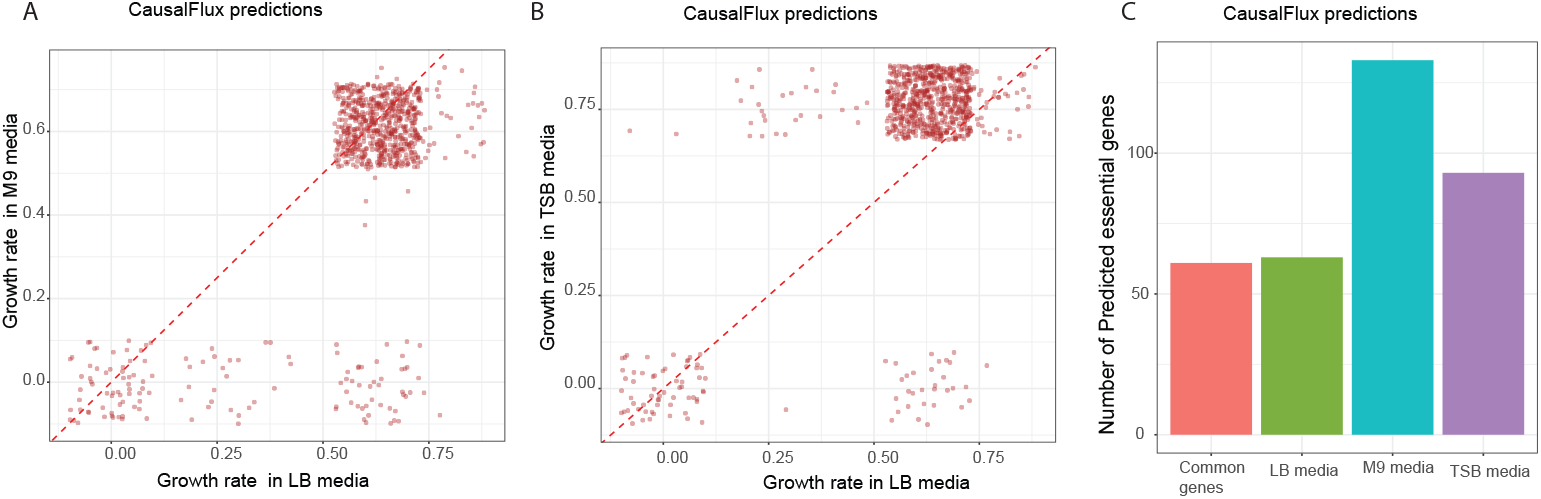
CausalFlux predictions under different media conditions: (A, B) Each is a scatter plot (with jitter 0.1 units) of predicted growth rate (biomass) under the two indicated media conditions. (C) Bar plot showing the number of predicted essential genes (predicted growth rate of zero) under the indicated media conditions. Common genes (orange) represent the number of predicted essential genes shared across all media conditions.

In 70 single-gene KO cases, growth was observed in LB but not in M9 predictions, indicating media-specific essentiality (see Table S3 for the gene list). Similarly, in 45 knockout cases, growth was observed in TSB but not in M9 predictions. Many of these genes, including *LeuD, LeuB, TrpA, TrpC, ArgH*, and *ArgC*, are involved in amino acid biosynthesis pathways such as leucine, tryptophan, and arginine synthesis. This result is expected because M9 minimal medium lacks amino acids; consequently, knockouts in these biosynthetic pathways lead to zero-growth conditions.

## Discussion

Integrating gene regulatory networks (GRNs) and genome-scale metabolic models (GSMMs) while accounting for bidirectional feedback mechanisms is crucial for accurately modeling and understanding the complex interplay within biological systems. In this work, we introduce CausalFlux, a novel methodology that integrates GRNs and GSMMs to predict reaction fluxes under gene KO perturbations. CausalFlux operates through an iterative process that combines causal surgery on the GRN to simulate the impact of gene KOs with constraint-based analysis of the GSMM to predict the resulting metabolic fluxes. We evaluated the performance of CausalFlux using three testbed datasets, which enabled direct comparison of predicted steady-state fluxes with ground-truth values obtained from ODE-based simulations. Across most evaluation scenarios, CausalFlux demonstrated a clear advantage over TRIMER in capturing trends in biomass fluxes across different testbed models, and this performance advantage was consistently observed under varying conditions, such as different exchange rates. These results underscore the importance of incorporating feedback mechanisms into integrated modeling frameworks. We further applied CausalFlux to predict growth versus no-growth phenotypes in *E. coli* across 798 single-gene KO cases, achieving a balanced accuracy of 0.79, which again exceeded that of TRIMER. Additionally, we predicted gene essentiality for these 798 genes under multiple growth media conditions, including TSB and M9 minimal media. Together, these analyses demonstrate the potential applicability of our algorithm. Overall, our results suggest that CausalFlux effectively captures reaction flux trends (particularly qualitative trends such as significant increases or decreases in reaction fluxes or the ability to distinguish zero versus nonzero growth outcomes) under gene KO perturbations.

In testbed models, we clearly see that Causalflux outperforms other state-of-the-art method like TRIMER in most of the KO conditions across all the testbed models. While the existing method GIMME exhibited comparable to or sometimes better performance than CausalFlux in predicting biomass reaction fluxes of testbed models, it is important to note that GIMME requires extra information in the form of gene expression data specific to the KO condition. In contrast, CausalFlux achieves similar predictive power without relying on such condition-specific data, and only needs expression data from a broad set of conditions to learn the initial Bayesian GRN, making it more broadly applicable.

Several studies have emphasised the role of metabolites in regulating gene expression in microorganisms, especially *E. coli*. Geng and Jiang, in their work, mention cAMP, a well-known metabolite to regulate the *crp* gene, which is considered to be a global regulator in *E. coli* [20]. Weeramange et al. discuss how the *cra* gene in *E. coli* is regulated through the binding of fructose-1-phosphate, and note that this regulatory mechanism has been experimentally validated in earlier studies [21]. Cui et al. in their study state the effect of the metabolite leucine in regulating the *lrp* gene in *E. coli* [22]. Kumar et al. curated 77 metabolite-to-gene interactions involving 66 unique metabolites and 57 unique genes [12]. Given the significance of such metabolic feedback mechanisms as mentioned in the earlier studies, we incorporate these metabolite-to-gene feedback interactions into our CausalFlux algorithm. Using this information about the feedback edges, we performed 798 single-gene KOs in *E. coli* to identify essential genes. A closer inspection of this real-world analysis of gene essentiality reveals that incorporating metabolic feedback into our algorithm results in performance improvements at each iteration, with the F1 score increasing over time. The ablation study that we performed, in which metabolite-to-gene edges were removed before running CausalFlux (on the 798 single-gene KOs), demonstrated the importance of modeling feedback, as performance declined after edge removal. We could see the decrease in the F1 scores of the models where the edges from metabolite to gene, like *crp* or the top 10 metabolic feedback genes, were removed. Some of the non-essential genes like *astC, cysA, argD*, etc., were misclassified as essential genes (i.e., the biomass after the gene KO was zero).

There are some caveats to our methodology. CausalFlux currently provides qualitative predictions of reaction fluxes. This limitation arises from the lack of kinetic parameters, which are essential for precise quantitative modeling of the integrated system dynamics. Also, our algorithm can predict only single-gene KOs and at present cannot handle multiple-gene KOs. In addition to comparing our method with TRIMER, we also evaluated our approach with earlier approaches like rFBA and PROM [5, 8]. Since rFBA is significantly older and no functional implementation is currently available, a direct evaluation on predicting the essentiality of the 798 genes in *E. coli* was not feasible. However, PROM benchmarks itself with rFBA and provides the results for both methods on a separate dataset and gene KO conditions. We therefore assessed our method on the same dataset and gene KO cases as performed by Chandrasekaran et al., and compared our findings with the published performance values for rFBA and PROM to allow a fair comparison. Our findings demonstrate that CausalFlux outperforms rFBA at all thresholds used to classify gene KOs as essential (no-growth cases) or non-essential (growth cases) on the dataset utilized by Chandrasekaran et al. Relative to PROM, our method has comparable performance in one of the three thresholds, while PROM remains better in others. These comparisons show that CausalFlux remains competitive with both recent and earlier methods.

Nevertheless, CausalFlux’s ability to capture bidirectional interactions between the gene regulatory and metabolic networks offers a more complete understanding of cellular behaviour and has the potential to significantly impact fields such as metabolic engineering, drug discovery, and systems biology. In future, this methodology can be potentially applied to other model organisms, such as yeast and *B. subtilis* to understand the bidirectional feedback.

## Materials and Methods

### Background on separate GSMM and GRN models

GSMM models a network of interconnected metabolic reactions in a cellular system using six components: set of all metabolic reactions in the model (*n* reactions), set of all metabolites participating in these reactions (*m* metabolites), stoichiometric matrix (***S*** ∈ ℝ^*m×n*^) representing the metabolite substrates and products of the reactions, vector of lower bounds on reaction fluxes (***v***_***lb***_ ∈ ℝ^*n*^), vector of upper bounds on reaction fluxes (***v***_***ub***_ ∈ ℝ^*n*^), and set of gene-protein-reaction (*GPR*) relations. A GPR relation specifies the activity of a metabolic reaction using Boolean-logic-based rules (specifically, a Boolean expression composed of “AND/OR” operators acting on the expression states of genes, such as enzyme-coding genes, that regulate the reaction). Steady-state fluxes of reactions in a GSMM can be predicted using well-known methods such as Flux Balance Analysis (FBA) [23] and Flux Variability Analysis (FVA) [24].

On the other hand, GRN model of transcriptional regulation is a directed probabilistic graphical model, also known as a Bayesian network [25]. This Bayesian or probabilistic GRN model has two components: (i) [GRN structure] a directed graph whose nodes are all genes in the system (including transcription factors and target genes), and directed edges represent regulatory influences between them, such as the control of a target gene by a transcription factor; and (ii) [probabilistic model] a set of parameters that describe the probability distributions governing the binary expression states of all genes in the network (with each node/gene associated with a binary-valued random variable that is 0 for inactive and 1 for active state of the gene). Being a Bayesian network, the graph is directed and acyclic, and the joint distribution of its random variables factorizes as the product of the conditional distribution of each gene given its parents in the GRN. That is, 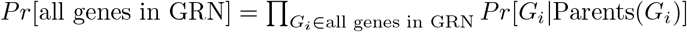. More formally, if *G*_1_, …, *G*_*p*_ denote the random variables corresponding to all *p* genes in the GRN, and *g*_*k*_ ∈ {0, 1} the binary state of *G*_*k*_,

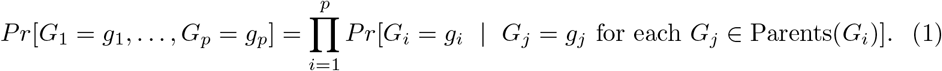

Hence, the parameters of the local conditional distributions (learnt from data) suffice to specify the global joint distribution.

### CausalFlux method

CausalFlux integrates the separate models of the two cellular subsystems described above (GSMM for metabolism and GRN for transcriptional/gene regulation) by crucially utilizing the bidirectional feedback interactions between them. The goal is to simulate the coupled GRN and GSMM models, and thereby predict the steady-state flux of reactions in the GSMM under this two-way feedback. The GRN-to-GSMM feedback is specified using the GPR rules, and the reciprocal GSMM-to-GRN feedback is specified by a set of feedback metabolites (***M***_***fb***_ ⊆ all metabolites in GSMM) and the genes they regulate. A key contribution of CausalFlux is to realize this reciprocal feedback using the concept of “causal surgery” on the Bayesian GRN and of “sink reactions” added to the GSMM, both of which are elaborated in this section as part of the overall workflow.

Given GRN, GSMM, and feedback information (GPR rules and genes regulated by ***M***_***fb***_) as the primary inputs, our overall CausalFlux method consists of a one-time initialization procedure, followed by an integration step that is executed iteratively until convergence (see Fig. 1C and the pseudocode in Algorithm 1 below). To complete the specification of our CausalFlux method, we expand on the initialization procedure, the integration step, and the iterative execution and convergence in the following three subsections.

#### Algorithm 1 CausalFlux

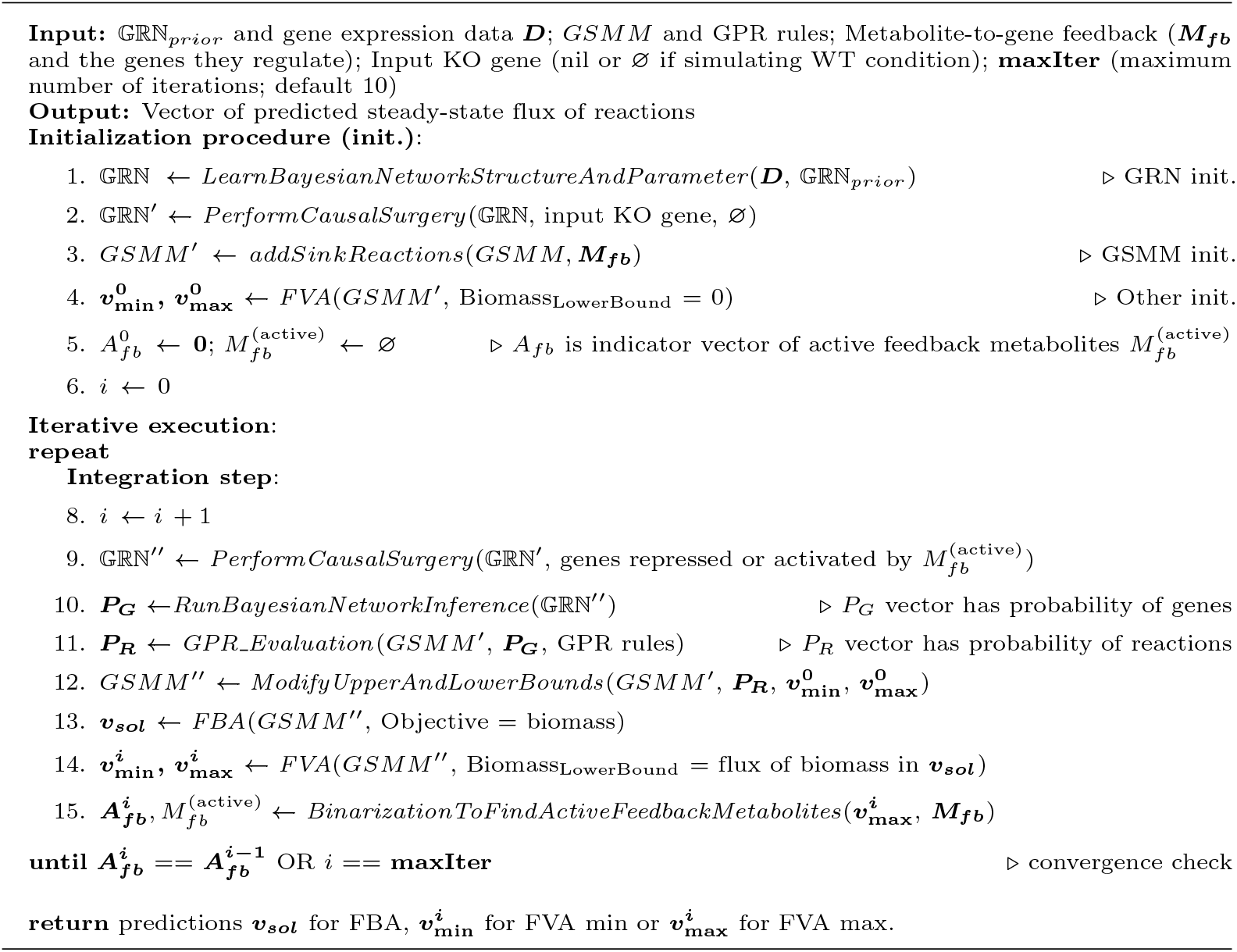

### CausalFlux: Intialization procedure

#### GRN Initialization (Bayesian network learning and initial surgery)

Given a genome-wide gene expression (transcriptomic) dataset and a prior GRN structure as input, we learn the Bayesian network structure and associated parameters of the Bayesian GRN using the R package *bnlearn* [26]. In detail, we used the hill-climbing heuristic in *bnlearn* for Bayesian network structure learning from the binarized gene expression dataset and the prior edges present between the genes (provided as a “white list” of edges to be required to be present in the learnt structure). Binarization of gene expression data is done in a dataset-specific fashion, and details on the same are given in the Methods sections below on application of CausalFlux to different datasets/models. For structure learning, Bayesian Dirichlet equivalent was used as the score/objective function, and other hyperparameters were set at default values, except for the maximum number of parents and maximum iterations, which were set at 5 and 100, respectively. The learnt structure will contain all the prior edges (“white list”) and possibly also additional edges supported by the data. With this learnt structure, we perform a final maximum likelihood-based parameter estimation to learn the conditional probability distribution of each gene in the GRN. The procedure explained here corresponds to *LearnBayesianNetworkStructureAndParameter* step in Algorithm 1.

If an input KO gene denoted *A* is given, then fluxes must be predicted under this causal perturbation. This initial perturbation is realized via causal surgery of the Bayesian GRN that removes edges incoming to *A* from its parents and forces the inactive (0) state on *A* with *do*(*A* = 0) operation (see below for more details on causal surgery).

#### GSMM initialization (sink reactions and flux bounds)

Standard constraint-based modeling methods like FBA or FVA can be applied on the GSMM to infer reaction fluxes, but metabolite concentration or activity levels cannot be directly inferred. Still, CausalFlux needs to know the activity of feedback metabolites ***M***_***fb***_ in the GSMM to capture their impact on the GRN. To achieve this, we add a sink reaction for each feedback metabolite to the GSMM. This maps to the *addSinkReactions* step in Algorithm 1. A sink reaction is an exchange reaction with a zero lower flux bound; therefore, it allows secretion but prevents uptake of the corresponding metabolite by the system. If an exchange reaction is already present for a metabolite *µ* ∈ ***M***_***fb***_, then we do not add an extra sink reaction. We simply force this reaction’s lower flux bound to zero, provided this metabolite is not an essential metabolite that needs to be uptaken by the system to grow. If it is an essential metabolite, then the lower bound is kept at a minimal non-zero value (that is specified in the Methods sections below on application of CausalFlux to different datasets/models). For convenience, we refer to all exchange reactions involving the feedback metabolites as sink reactions hereafter. The fluxes through these sink reactions are used to determine the activity of the feedback metabolites. The upper bounds for these sink reactions are kept in such a way that the optimal biomass reaction flux obtained by FBA does not change. Details on the bounds of sink reactions are given in the Methods sections below on application of CausalFlux to different datasets/models.

After all necessary sink reactions are added, we perform FVA on the GSMM to obtain the minimum 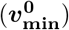 and maximum 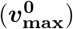 possible flux through each reaction (with no restriction on the achievable biomass, other than the biomass being at least 0). These two flux vectors are later used to modify the lower and upper bounds of the metabolic model in the “Integration” step.

### CausalFlux: Integration step

#### Causal surgery to mutilate GRN and perform inference

Causal surgery on a random variable in a (causal) Bayesian network is an ideal intervention on the variable, in which its value alone is externally fixed and its dependence on its parent variables is removed [25]. More generally, any intervention on a node in a causal graph is referred to as causal surgery [27]. Such an intervention severs the incoming edges from the parents of the target node, resulting in a mutilated graph, and fixes the conditional probability of the intervened node to a specified value. Mathematically, this operation is represented using the “do-operator”, which defines the probability distribution of a target variable when a conditioning variable is externally set to a particular value. A key contribution of the CausalFlux framework is its ability to simulate feedback effects of metabolites on genes through causal surgery on appropriate genes. This procedure is formalized in the *PerformCausalSurgery* step of Algorithm 1, which takes as input the GRN along with sets of repressed and active genes. Using the “do-operator”, genes in the repressed set are fixed to 0, while genes in the active set are fixed to 1, after removing the corresponding parental edges in the GRN. Importantly, causal surgeries on multiple genes can be performed in any order without affecting the resulting GRN structure.

To illustrate this process, consider Fig. 1B. Prior to causal surgery, the conditional probability of gene “D” depends on gene “C”, expressed as *P*(*D* = 1 | *C* = 1), where the state of “C” itself depends on its parent genes “A” and “B”. When a metabolite (M2) activates gene “C”, the “do-operator” is applied to set *C* = 1, yielding *P*(*D* = 1 | *do*(*C* = 1)). This intervention removes the incoming edges from “A” and “B” to “C” and fixes the activity of “C” to 1.

Once the causal surgery above is done, we perform inference on the mutilated Bayesian network to infer the probability of each gene (collected in the vector *P*_*G*_ across all genes) – this process corresponds to the *RunBayesianNetworkInference* step of the Algorithm 1. To clarify, for each gene, the above probability is actually the conditional probability of the gene in the original Bayesian network given all the do(.) operations on the affected genes (which includes the input KO gene as well as the genes controlled by active feedback metabolites; note that the effect of input KO gene is also captured via causal surgery as mentioned earlier, wherein the edges between the input KO gene and its parents are severed and the input KO gene is forced to activity 0). This is done using the *cpquery()* function from the *bnlearn* package in R [26].

#### Bounds modification and constraint-based analysis

The conditional probability of genes as calculated above is next used to obtain the conditional probability of the activity of all metabolic reactions 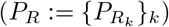 using the corresponding GPR rules. This is performed by the *GPR Evaluation* step of Algorithm 1, which in turn uses the *GPReval()* function from the COBRA Toolbox. In detail, this function analyzes the *k*-th metabolic reaction denoted *R*_*k*_ (for all *k*) as follows. The conditional probability 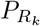 of reaction *R*_*k*_ is obtained by taking the maximum of the conditional probability of genes in an OR rule in the GPR of *R*_*k*_, and the minimum of conditional probability of genes in an AND rule in the GPR of *R*_*k*_; for complex GPR rules with multiple OR and AND operators, *GPReval()* function uses standard operator precedence conventions (i.e., prioritizes operators in the order: parentheses, AND, and OR) [28].

Once the above probabilities are computed, the bounds of the *k*-th metabolic reaction is updated (by the *ModifyUpperAndLowerBounds* component of Algorithm 1) using the equations below.

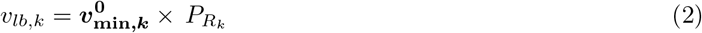

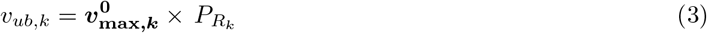

The updated bounds of all reactions are incorporated into the metabolic model to perform FVA and FBA.

#### Determination of activities of metabolites

By constructing a binarized vector with values of 1 or 0, the activity of metabolites in ***M***_***fb***_ is determined to be active or inactive, respectively. In the case of CausalFlux, the following Eq 4 describe the binarization procedure,

For any *µ* ∈ ***M***_***fb***_, we’ve

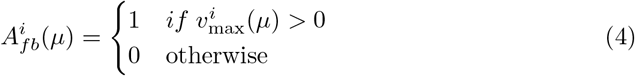

Here, 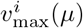 refers to the maximum flux obtained from FVA, passing through the sink reaction corresponding to feedback metabolite *µ* in the *i*^*th*^ iteration of CausalFlux. The above procedure is captured within the *BinarizationToFindActiveFeedbackMetabolites* component of Algorithm 1.

### CausalFlux: Iterative execution and convergence

The integration step described above corresponds to one iteration, and we repeatedly execute the integration step until convergence.

#### Checking convergence

The *A*_*fb*_ vector indicating the activity status of ***M***_***fb***_ in the *i*^*th*^ vs. (*i* − 1)^*th*^ iterations are compared. If they are identical, the algorithm converges. When these vectors are not identical, the active metabolites in ***M***_***fb***_ are incorporated back to the GRN to perform causal surgery on the genes regulated by the active metabolites. This iteration continues until the *A*_*fb*_ vectors obtained between the *i*^*th*^ and (*i* − 1)^*th*^ iterations are identical or till the iteration reaches the pre-defined maximum number of iterations.

#### Predictions

To determine the final biomass after the algorithm terminates, the following procedure was applied. If the algorithm terminated before reaching the maximum number of iterations, the algorithm has converged and hence the biomass value from that iteration was used as the final value. If the algorithm reached the maximum iteration limit, two scenarios were possible: the algorithm has either converged or stopped due to the iteration limit. To distinguish these cases, we examined the fluxes of the feedback metabolite reactions (exchange and sink). In case of convergence, the biomass from the last iteration was taken as the final biomass. In case of non-convergence (e.g., if oscillations were present in the exchange/sink fluxes), the average biomass flux from the last two iterations was taken as the final biomass.

### Testbed Model (TM) benchmarks

To evaluate CausalFlux’s predictions, we consider three simplified TMs of integrated GRN-GSMM systems (labeled TM1, TM2, and TM3). A testbed model is specified using its topology (network structure), and a system of ODEs based on this topology that govern the dynamics of the coupled GRN-GSMM system. Simulating these ODEs yields ground-truth steady-state fluxes of reactions in the testbed model, which can then be compared to steady-state fluxes predicted by CausalFlux applied on the testbed model.

### Topology of TMs

A testbed model consists of a small-scale GRN-like and GSMM-like component. The network motifs prominent in biological systems served as the basis for constructing the GRN components in the testbed models. We incorporated some well-known network motifs, such as feedforward and feedback loops, single-input modules, and dense overlapping regulons, into our testbed models due to their prevalence in the transcription networks of biological systems [29]. The small-scale GSMM comprises a few reactions, which operate on metabolites, and are catalyzed by enzymes (as per the GPR rules based on enzyme-encoding genes). For simplicity, we formulate all these metabolic reactions as irreversible reactions. In all TMs, we treat reaction *R*1 as the exchange (nutrient import) reaction. We specify the biomass reaction to be the reaction R4 in TM1, R11 in TM2, and R8 in TM3. This specification is required when three methods, CausalFlux, TRIMER, and GIMME, are applied on the testbed models to predict fluxes. The topology of TM1 is depicted in Fig. 2A (and of TM2 and TM3 in S1 Fig), and the components of TMs (including metabolic reactions and gene-gene interactions) are in Table S1.

### ODE representation of TMs

The dynamics of the interacting molecules in TMs is represented using ODE equations. The system of ODEs representing the different reactions in the TMs follows Michaelis–Menten kinetics [30]. The following set of parameters are used to describe reactions in the TMs: Michaelis–Menten kinetics maximum reaction velocity, *V*_*max*_ (mM/s); Michaelis–Menten kinetics maximum substrate concentration, *K*_*m*_ (mM); sink constant required for outflux reactions, *K*_*s*_ (mM/s); and time constant, T (s). Taking TM1 as an example, this section describes the different reaction types in TM1 and the ODEs that govern their kinetics; TM2 and TM3 can be described similarly using their corresponding ODEs (listed in Suppl Section A.1). A TM has five types of reactions:

1. Nutrient import reaction (R1): All TMs have rate of nutrient uptake proportional to the biomass reaction and the negative of the intracellular concentration of the nutrient. The maximum nutrient uptake rate was fixed to 3.2 mM/s [31]. The nutrient uptake reaction for TM1 is given by:

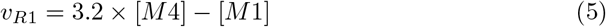
2. Metabolite export reactions (R6 and R7): The rate of these reactions is directly proportional to the concentration of the metabolites. For example, the rate of reaction 6 is given by:

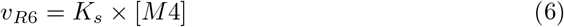
3. Enzymatic reactions (R2, R3, R4, and R5): All enzymatic reactions are modeled using Michaelis–Menten kinetics [30] with its two associated parameters: maximum reaction rate, *V*_max_ and Michaelis–Menten constant, *K*_*m*_. *V*_max_ is modified to incorporate the GPR rules. To illustrate this for reactions R2 and R3, please see the equations below.

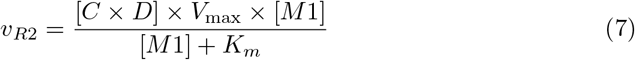

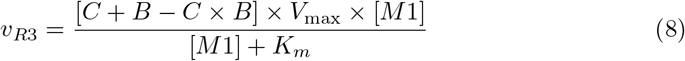
4. Gene-Gene interaction reactions: Gene-gene interactions are captured using the normalised Hill differential equations [32]. “*F*_*act*_(gene)” here represents the fractional activation of a gene that follows normalised Hill kinetics. Gene B is represented by:

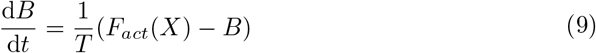
5. Metabolite–Gene interaction reactions: “*F*_*act*_(*metabolite*)” here represents the fractional activation of a metabolite, indicating 1 if the species is greater than 50% of concentrations and 0 otherwise. Gene X from TM1 is represented by:

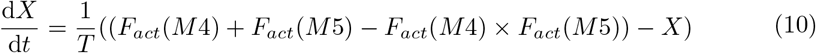

Here, [*M*4], [*M*5] and [*M*1] refer to the concentrations of the metabolites M4, M5 and M1, respectively. Also, *C, D*, and *B* refer to the fractional activation values of the genes C, D, and B, respectively.

### Simulation of ODE models to generate ground-truth steady-state fluxes and gene expression data

Each of the three TMs we consider can be simulated under WT or different gene KO conditions, and under three exchange rates. In total, this yields 39 TM scenarios/cases to simulate as shown in Fig. 2E. For each of the 39 TM cases (specified by the TM, WT/gene-KO condition, and exchange rate), we simulate the corresponding system of ODEs under 100 distinct parameter configurations. These parameter configurations were constructed from parameter ranges of the ODE parameters, *V*_max_, *K*_*m*_, *K*_*s*_, and *T* shown in Table S2, which were in turn based on parameter ranges reported in an earlier study by Lee et al. [31] on an ODE-based modeling framework for metabolic, regulatory, and signaling processes.

Given any of the 39 TM cases, we now elaborate on how we simulate the TM case and generate relevant data. For each of the 100 distinct parameter configurations of the given TM case, we simulate the corresponding ODE system for 10,000 time points, and record the reaction fluxes and gene expression data from the last time point as the data pertaining to steady-state behavior. This results in 100 sets of steady-state data (on reaction fluxes and gene expression) pertaining to the different parameter configurations. The median of these 100 steady-state flux values for each reaction is taken as the ground-truth steady-state reaction fluxes for the given TM case. In contrast, all 100 simulated steady-state gene expression values were retained without aggregation, resulting in 100 expression instances per gene for the TM case. These datasets can then be used to benchmark methods like CausalFlux or TRIMER. The rationale for aggregating steady-state flux values is due to the constraint-based GSMM that underlies the benchmarked methods, like CausalFlux or TRIMER in detail, the GSMM components of these methods do not rely on any knowledge of the kinetic parameters and are hence tuned to predict a steady-state behavior (reaction fluxes) that is only representative of the actual steady-state behavior. We take this representative to be the median of a set of possible steady-state behaviors for this benchmark. The rationale for not aggregating gene expression instances is to enable learning of the GRN component of these methods from multi-sample gene expression data.

We provide some additional information on the different exchange rate settings and related analyses. The ODE systems of TMs were simulated under WT/gene-KO condition across three exchange rates, the default 3.2 mM/s rate, and two additional scaled rates of 320 and 3200 mM/s. To simulate scaled exchange rates, parameters *V*_max_ and *K*_*s*_ corresponding to the 3.2 mM/s exchange rate were each scaled proportionally by 100-fold and 1000-fold for the 320 and 3200 exchange rates, respectively. When applying CausalFlux, TRIMER, or GIMME, the 100 gene expression instances generated for a given TM, condition, and exchange rate were treated as independent samples. Refer to S2A Fig for a schematic overview of the simulation pipeline.

### Application of methods on TM benchmarks

#### Application of CausalFlux on TM benchmarks

We used CausalFlux to predict steady-state fluxes for all reactions in testbed models, TM1, TM2, and TM3, under WT and KO conditions across three different exchange rates (a total of 39 TM cases; see Fig. 2E). To apply CausalFlux on each such TM case (i.e., a TM under a given exchange rate and a WT or gene-KO condition), the following inputs were used: **(1)** 𝔾ℝℕ_*prior*_ **and binarized gene expression data *D*:** The actual topology of the TM is provided as 𝔾ℝℕ_*prior*_ to CausalFlux (see Table S1). The continuous-valued gene expression data (obtained from simulating the system of ODEs of the TM for the WT condition under the given exchange rate) is binarized (converted to 0/1 data matrix ***D***) using k-means clustering (binarize.kMeans() function from the “BiTrinA” package [33] was used with k = 2). Please note that CausalFlux’s step *LearnBayesianNetworkStructureAndParameter* can learn some additional edges based on ***D***, besides including the edges provided in 𝔾ℝℕ_*prior*_. Additionally, note that the input ***D*** is always based on the WT simulated expression data, regardless of whether the TM case on which CausalFlux is applied specifies the WT or KO condition, in order to mimic real-world settings where expression data is more often available under WT than gene-KO conditions. **(2)** *GSMM* **and GPR rules:** The small-scale GSMM component of a particular testbed model is provided as input (see Table S1 for details on the GPR rules, reactions, and metabolites for each TM). The lower bound for reaction “R1” in the metabolic model is set to the exchange rate used in ODE simulations. The upper bounds of all the reactions were set at 10000; **(3) Metabolite-to-gene feedback:** Edges from metabolites to genes for each testbed model (see Table S1); and **(4) Input KO gene:** nil or ∅ for the WT, and a specific gene for the KO condition (refer to Fig. 2E for all KO genes, each of which were separately knocked out).

Once the GRN is obtained, the remaining components of the methodology are implemented as mentioned earlier. The steady-state fluxes of every WT/KO condition were predicted for every TM under every exchange rate. The correlation (*ρ*) between the actual and predicted fluxes (CausalFlux) was calculated for all the TMs under WT/KO cases. The maximum number of iterations for running CausalFlux on TMs was set at 10. While plotting the scatter plots between the *ρ* computed between actual and predicted fluxes across the various TM cases, “jitter” from “ggplot2” function was used to avoid overplotting of the points in Fig. 2[D-E] and Fig. 3[A-F].

#### Application of other methods on TM benchmarks

GIMME (from COBRA Toolbox) [28] and TRIMER were applied to the same 39 TM cases as CausalFlux to predict steady-state fluxes [2, 9]. Although these two studies differ significantly from CausalFlux in terms of input types and approach, we wanted to assess how well they performed on the TMs to estimate fluxes under WT or gene-KO conditions. For each TM case, the GRNs and binary gene expression data provided as input to CausalFlux is also provided as input to execute TRIMER. “sFBA” was used to predict the steady-state fluxes in TRIMER, since it was their best performing mode [9]. For each TM case, GIMME predicted fluxes using the average gene expression data simulated under the particular condition (WT or gene-KO condition) and exchange rate specified in the TM case.

### Application of methods on real-world datasets

#### Application of CausalFlux on *E. coli* to perform single-gene KO perturbations

We applied CausalFlux to perform single-gene KOs on 798 *E. coli* genes under LB (Luria–Bertani) media to predict cellular growth. Ground-truth growth rates for these knockout cases were obtained from the Keio collection, where zero measured growth indicates no growth and non-zero values indicate growth [13]. The 798 genes were selected because they had at least one edge in the GRN learnt/reconstructed from the *E. coli* expression data and were included in at least one GPR rule of *E. coli* ‘s metabolic model (see below). Growth predictions were inferred from CausalFlux-predicted biomass flux, with zero biomass indicating no growth and nonzero biomass indicating growth.

In this application of CausalFlux, the relevant inputs/datasets are as follows: **(1)** 𝔾ℝℕ_*prior*_ **and binarized gene expression data *D*:** Prior knowledge (𝔾ℝℕ_*prior*_) on gene-gene pairwise interactions was taken from RegulonDB v8.1 (3451 unique gene-gene pairs without any cycles), similar to TRIMER [34]. The binarized gene expression data for *E. coli* was taken from the TRIMER work, with the underlying raw expression data of 4189 genes in 2198 samples based on the EcoMac work [35]. As described by the TRIMER authors, the raw expression data were quantile-normalized and subsequently binarized using a quantile-based threshold reported as 0.33 (hence if a gene’s normalized expression in a given sample is less than or equal to the gene’s 33rd percentile value, it is considered OFF (0), and otherwise considered ON (1)). The resulting OFF/ON (0/1) binarized gene expression data is used for Bayesian network learning. The learnt 𝔾ℝℕ for *E. coli* consisted of 3551 unique edges over 1607 genes. **(2)** *GSMM* **and GPR rules:** We used iML1515 model for *E. coli*, consisting of 1877 metabolites, 2712 reactions, and 1516 genes [36]. The GPR rules are also taken from this model. The lower bound of essential exchange reactions for feedback metabolites was set at − 0.001, and the upper bound of the sink reactions for feedback metabolites was set at 10. The lower bound of *v*_*glucose*_ (glucose exchange rate) and *v*_*oxygen*_ (oxygen exchange rate) in the GSMM was set at − 8.5 mm/gDCW/hr and − 14.5 mm/gDCW/hr respectively, as described in TRIMER for aerobic conditions [9]. To simulate the LB media condition, we modified the bounds for some of the exchange reactions that are present in the media, as done in the work by Zimmermann et al. [37]. **(3) Metabolite-to-gene feedback:** 77 unique edges between 66 feedback metabolites ***M***_***fb***_ and 57 (transcription factor) genes in *E. coli* taken from Kumar et al. [12]. **(4) Input KO gene:** We applied CausalFlux to separately perform each of the 798 single-gene KOs. The Keio collection has the growth rate experimentally measured for 3985 KO strains in *E. coli* under LB media [13]. Additionally, we relied on the EcoCyc database [38] to determine whether a gene is essential vs. non-essential for *E. coli* grown in LB media (note that an essential gene is one whose KO leads to no growth, and is hence necessary for growth).

#### Application of other methods on *E. coli* to perform single-gene KO perturbations

On the *E. coli* dataset covering 798 single-gene KO perturbations, the TRIMER algorithm was applied using the same binarized gene expression data, the same set of prior edges, and the same iML1515 metabolic model as used for CausalFlux. The glucose and oxygen exchange rates, as well as the LB medium setup, were also identical to those used in the CausalFlux application. Again, steady-state fluxes in TRIMER were predicted using the sFBA mode, as it was reported to be the best-performing configuration by TRIMER authors.

We did not apply GIMME to the 798 single-gene KO perturbations because it requires gene expression data for each KO condition, which is difficult to obtain under LB medium. Moreover, since our objective is to predict gene essentiality in *E. coli*, acquiring steady-state gene expression data for essential genes is inherently troublesome, as their KO would result in cell death.

#### Application of CausalFlux on *E. coli* to perform single-gene KO perturbations under different media conditions

We modified the lower and upper bounds for exchange reactions corresponding to M9 and TSB media in the iML1515 *E. coli* model, with values taken from Zimmermann et al. [37]. After modifying the GSMM, CausalFlux was applied as described in the previous section (on its application on *E. coli* to perform single-gene KO perturbations under the LB media).

#### Application of CausalFlux on rFBA/PROM *E. coli* datasets to perform 15 gene KOs across 100 media conditions

CausalFlux was applied separately to the rFBA and PROM datasets to predict biomass (growth rate) for 15 gene knockouts (KOs) across 100 media conditions. These 100 media conditions comprised 58 carbon sources, 31 nitrogen sources, and 11 double-nitrogen sources.

For the rFBA datasets, the GRN information and feedback metabolite data were extracted from Covert et al.’s work [5]. The iJR904 metabolic model was used for this analysis. Since rFBA itself does not require gene expression data but CausalFlux does, the gene expression data were taken from Chandrasekaran et al.’s work [8].

For the PROM datasets, both the GRN information and the gene expression data were obtained from Chandrasekaran et al.’s work. As the PROM framework does not include metabolic feedback, the feedback information used in CausalFlux was kept consistent with that used in the rFBA analysis. The iJR904 metabolic model was also used for this case.

Chandrasekaran et al. presented results using the iJR904 model of *E. coli* and provided rFBA predictions on the same model. The ground truth labels (lethal/non-lethal cases), along with the rFBA and PROM predictions, were obtained from their work. Covert et al. also provided the exchange reaction bounds in the iJR904 model required to simulate the 100 media conditions.

### Applying CausalFlux following ablation of metabolite-to-gene feedback edges

The metabolite-to-gene feedback interactions in *E. coli* considered in this work involved 54 unique genes, referred to as metabolic feedback genes. We define the influence of any such metabolic feedback gene (on the single-gene KOs) as the number of single-KO genes that are in the 0-hop or 1-hop neighbourhood of that gene in the GRN. The metabolic feedback genes can then be ranked based on this influence measure. We conducted five separate ablation studies in which the feedback edges from metabolite(s) to gene(s) were removed and the change in prediction performance noted. Specifically, in the first three studies, we ablated feedback edges to *crp, fur*, and *cra*, which are the metabolic feedback genes with the highest, second-highest, and third-highest influence (on the single-gene KOs) respectively. In the final two ablation studies, we considered a set of 10 (and separately 40) metabolic feedback genes with the most influence on the single-KO genes (with the latter representing approximately 75% of the total 54 genes) and removed the incoming feedback edges from metabolites to all genes in the set. After performing any of the above ablations of metabolite-to-gene edges, the CausalFlux algorithm was applied to predict the biomass for each of the 798 single-gene KO conditions, and these predictions compared to the ones obtained before ablation.

## Supporting information

Supplementary Information (Supplementary Methods/Figures/Tables/DataFiles)

## Supporting information

**S1 Fig. Topology of TM2 and TM3:** The integrated GRN+MM network for TM2 and TM3 are shown.

**S2 Fig. Framework for simulating ground-truth fluxes/gene expression for a particular TM case; and evaluating CausalFlux, TRIMER, and GIMME on this TM case:** Consider a system of ODEs representing a given TM case, i.e., a TM (integrated small-scale GRN+GSMM testbed system with two-way feedback) under a specific WT/KO condition and a particular exchange rate. (A) For each of the 100 parameter configurations shown, we simulate this ODE system for 10,000 time points to eventually obtain steady-state data (reaction fluxes and gene expression) matrix for the given TM case. (B) Different methods (CausalFlux, TRIMER, and GIMME) are applied on the resulting steady-state data to generate predicted fluxes. The median value of steady-state fluxes across the 100 parameter configurations is taken as the actual or ground-truth fluxes. The correlation (*ρ*) between the predicted and ground-truth fluxes can be used to evaluate the different methods.

**S3 Fig. Comparison of GIMME variants (FVA max, FVA min and FBA) with actual in TMs:** Scatter plot of the correlation (*ρ*) computed between the actual and GIMME (FVA max) for the 39 TM cases is compared with *ρ* computed between actual and GIMME (FBA), (B) actual and GIMME (FVA min).

**S4 Fig. Comparison of CausalFlux iteration 1 vs 2 for 39 TM cases:** Scatter plot (with jitter of 0.05 units) between the *ρ* computed between actual and CausalFlux predicted fluxes at iteration 1 versus iteration 2 for the 39 TM cases.

**S5 Fig. Exploring the role of metabolic feedback through ablation studies:** Scatter plots (with jitter of 0.05 units) for ablation studies done on (A)*fur*, (B) *cra*, (C) Metabolic feedback to 40 metabolic feedback genes with the most influence on single-gene KOs were removed.

**S6 Fig. Threshold of** 5% **of WT is used to define essential/non-essential cases**: (A) Scatter plot (with jitter 1.75 units) between the accuracy computed between CausalFlux predictions and Actual vs. the accuracy between rFBA predictions and Actual for the 100 media conditions. (B) Scatter plot (with jitter 1.75 units) between the accuracy computed between CausalFlux predictions and Actual vs. the accuracy between PROM predictions and Actual for the 100 media conditions. (C) Performance comparisons of CausalFlux and rFBA for the various Carbon sources, Nitrogen sources and Double-Nitrogen sources and overall 100 media conditions. (D) Performance comparisons of CausalFlux and PROM for the various Carbon sources, Nitrogen sources and Double-Nitrogen sources and overall 100 media conditions. (E) Comparison of classification metrics (TP: true positives [essential], FN: false negatives, FP: false positives, TN: true negatives [non-essential]) for CausalFlux applied to the rFBA and PROM datasets, along with Random Classifier, rFBA, and PROM predictions across 1500 data points (100 media conditions *×* 15 gene KOs).

**S7 Fig. Threshold of** 50% **of WT is used to define essential/non-essential cases**: (A) Scatter plot (with jitter 1.75 units) between the accuracy computed between CausalFlux predictions and Actual vs. the accuracy between rFBA predictions and Actual for the 100 media conditions. (B) Scatter plot (with jitter 1.75 units) between the accuracy computed between CausalFlux predictions and Actual vs. the accuracy between PROM predictions and Actual for the 100 media conditions. (C) Performance comparisons of CausalFlux and rFBA for the various Carbon sources, Nitrogen sources and Double-Nitrogen sources and overall 100 media conditions. (D) Performance comparisons of CausalFlux and PROM for the various Carbon sources, Nitrogen sources and Double-Nitrogen sources and overall 100 media conditions. (E) Comparison of classification metrics (TP: true positives [essential], FN: false negatives, FP: false positives, TN: true negatives [non-essential]) for CausalFlux applied to the rFBA and PROM datasets, along with Random Classifier, rFBA, and PROM predictions across 1500 data points (100 media conditions *×* 15 gene KOs).

**Table S1** For each TM, the genes, metabolites, and reactions in the TM are provided, which are in turn used to define and simulate a system of ODEs corresponding to the TM.

**Table S2** The range of parameter values used for simulating the ground truth data from the ODE equations. The parameter range shown here are for 3.2 exchange rate. *V*_*max*_ and *K*_*s*_ were scaled proportionally by 100-fold and 1000-fold for the 320 and 3200 exchange rates.

**Table S3** List of genes that vary under different media conditions when single-gene KOs are performed on *E. coli*.

**Table S4** For each metabolic feedback gene, this table shows its influence, i.e., the number of its 1-hop neighbors in the GRN that overlap with the set of 798 single-gene KOs in the *E. coli* dataset.

**Table S5** Performance metrics of the original model and 5 different ablation models for predicting essentiality of genes (i.e., growth vs. no-growth phenotype under single-gene KOs).

**Supplementary Data 1. Spearman’s** *ρ* **between actual and predicted fluxes for all the reactions in TMs**. Spearman correlation and its p-value computed between the actual and predicted fluxes (across all the methods) for the 39 cases of TMs are given in this file.

**Supplementary Data 2. Reactions with zero flux in the “gabT” and “serC” KO models with the metabolic feedback to “crp” removed**. This file lists the identifiers of the following reactions. A total of 123 reactions had zero flux, of which 27 had different GPR probabilities for the “gabT” KO model with metabolic feedback to “crp” removed. Similarly, a total of 144 reactions had zero flux, of which 29 had different GPR probabilities for the “serC” KO model with metabolic feedback to “crp” removed.

## Acknowledgments

We thank members of our research group BIRDS (Bioinformatics and Integrative Data Science) and research center IBSE (Centre for Integrative Biology and Systems Medicine) at IIT Madras for their valuable inputs during the course of this work. This work was supported by the Wellcome Trust/DBT India Alliance Intermediate Fellowship Grant IA/I/17/2/503323 awarded to MN. We acknowledge the use of LLMs (Large Language Models, mainly ChatGPT, Gemini, and Grammarly) for copy-editing purposes (checking spelling and grammar, and polishing certain sentences/phrases).

## Author contributions statement

**NS**: Conceptualization, Data Curation, Methodology, Software, Validation, Formal analysis, Writing - Original Draft, Writing - Review & Editing, Visualization. **SPK**: Conceptualization, Methodology, Software, Validation, Writing - Review & Editing, Visualization. **RR**: Supervision, Conceptualization, Visualization. **NPB**: Supervision, Conceptualization, Writing - Review & Editing, Visualization. **MN**: Supervision, Conceptualization, Methodology, Writing - Review & Editing, Visualization, Resources, Funding acquisition.

## Notes

https://github.com/BIRDSgroup/CausalFlux

### Competing Interest Statement

The authors have declared no competing interest.

