## Supplementary Information (Supplementary Methods/Figures/Tables/DataFiles) for "Modeling and dissecting bidirectional feedback in gene-metabolite systems using the CausalFlux method"

**B Supplementary Figures** **6**

**C Supplementary Tables** **13**

**D Supplementary Data Files** **17**

### A Supplementary Methods

#### A.1 Ordinary Differential Equations (ODEs) for the Testbed Models

##### A.1.1 Testbed model-1

**Reaction rates for every metabolic reaction in the system**

$$
\begin{aligned}
v_{R1} &= \mathsf{Exchange\_rate} \times M8 - M1 \\
v_{R2} &= ((B + C - (B \times C) \times M1 \times v_{max})/(M1 + K_m) \\
v_{R3} &= (J \times M1 * v_{max})/(M1 + K_m); \\
v_{R4} &= (K \times M4 * v_{max})/(M4 + K_m); \\
v_{R5} &= ((K + D - (K \times D)) \times M2 \times v_{max})/(M2 + K_m); \\
v_{R6} &= ((F + G - (F \times G)) \times M2 \times v_{max})/(M2 + K_m); \\
v_{R7} &= ((L + M - (L \times M)) \times v_{max} \times M3 \times M7)/(K_m + M3 + M7 + (M3 \times M7)); \\
v_{R8} &= ((H + G - (H \times G)) \times M5 \times v_{max})/(M5 + K_m); \\
v_{R9} &= (G \times M5 \times v_{max})/(M5 + K_m); \\
v_{R10} &= K_s \times M6 \\
v_{R11} &= K_s \times M8
\end{aligned}
\tag{S1}
$$

**Rate of change of metabolite's concentration in the system**

$$
\begin{aligned}
\frac{dM1}{dt} &= R1 - R2 - R3 \\
\frac{dM2}{dt} &= R2 + R4 - R5 - R6 \\
\frac{dM3}{dt} &= R5 - R7 \\
\frac{dM4}{dt} &= R3 - R4 \\
\frac{dM5}{dt} &= R6 - R8 - R9 \\
\frac{dM6}{dt} &= R9 - R10 \\
\frac{dM7}{dt} &= R8 - R7 \\
\frac{dM8}{dt} &= R7 - R11
\end{aligned}
\tag{S2}
$$

**Rate of change of activity of the genes in the system**

$$
\begin{aligned}
\frac{dA}{dt} &= T^{-1} \times (F_{act}(M8,1) - A) \\
\frac{dB}{dt} &= T^{-1} \times (F_{act}(A,0) - B) \\
\frac{dC}{dt} &= T^{-1} \times (F_{act}(A,0) - C) \\
\frac{dD}{dt} &= T^{-1} \times (F_{act}(A,0) - D) \\
\frac{dE}{dt} &= T^{-1} \times (Const - E) \\
\frac{dF}{dt} &= T^{-1} \times (F_{act}(E,0) - F) \\
\frac{dG}{dt} &= T^{-1} \times (F_{act}(E,0) - G) \\
\frac{dH}{dt} &= T^{-1} \times (F_{act}(E,0) - H) \\
\frac{dI}{dt} &= T^{-1} \times (Const - I) \\
\frac{dJ}{dt} &= T^{-1} \times (F_{act}(I,0) - J) \\
\frac{dK}{dt} &= T^{-1} \times (F_{act}(I,0) - K) \\
\frac{dX}{dt} &= T^{-1} \times (Const - X) \\
\frac{dL}{dt} &= T^{-1} \times (F_{act}(X,0) - L) \\
\frac{dM}{dt} &= T^{-1} \times (F_{act}(X,0) - M)
\end{aligned} \tag{S3}
$$

###### A.1.2 Testbed Model-2

**Reaction rates for every metabolic reaction in the system**

$$
\begin{aligned}
v_{R1} &= Exchange\_rate \times M4 - M1 \\
v_{R2} &= (C \times D \times M1 \times v_{max})/(M1 + K_m) \\
v_{R3} &= ((B + C - (B \times C)) \times M1 \times v_{max})/(M1 + K_m) \\
v_{R4} &= ((B + D - (B \times D)) \times M1 \times v_{max})/(M1 + K_m) \\
v_{R5} &= ((D + C - (D \times C)) \times M1 \times v_{max})/(M1 + K_m) \\
v_{R6} &= K_s \times M4 \\
v_{R7} &= K_s \times M5 \\
v_{R8} &= K_s \times M2
\end{aligned} \tag{S4}
$$

**Rate of change of metabolite's concentration in the system**

$$
\begin{aligned}
\frac{dM1}{dt} &= R1 - R2 - R3 \\
\frac{dM2}{dt} &= R2 - R4 - R8 \\
\frac{dM3}{dt} &= R3 - R5 \\
\frac{dM4}{dt} &= R4 - R6 \\
\frac{dM5}{dt} &= R4 - R5 - R7
\end{aligned} \tag{S5}
$$

**Rate of change of activity of the genes in the system**

$$
\begin{aligned}
\frac{dA}{dt} &= T^{-1} \times (F_{act}(M2,1) + F_{act}(Y,0) + F_{act}(Z,0) - (F_{act}(M2,1) \times F_{act}(Y,0) \times F_{act}(Z,0)) - A) \\
\frac{dB}{dt} &= T^{-1} \times (F_{act}(X,0) - B) \\
\frac{dC}{dt} &= T^{-1} \times ((F_{act}(A,0) + F_{act}(B,0) - (F_{act}(A,0) \times F_{act}(B,0))) - C) \\
\frac{dD}{dt} &= T^{-1} \times ((F_{act}(B,0) + F_{act}(E,0) - (F_{act}(B,0) \times F_{act}(E,0))) - D) \\
\frac{dE}{dt} &= T^{-1} \times (Const - E) \\
\frac{dY}{dt} &= T^{-1} \times (Const - Y) \\
\frac{dZ}{dt} &= T^{-1} \times (Const - Z) \\
\frac{dX}{dt} &= T^{-1} \times ((F_{act}(M4,1) + F_{act}(M5,1) - (F_{act}(M4,1) \times F_{act}(M5,1))) - X)
\end{aligned} \tag{S6}
$$

##### A.1.3 Testbed Model-3

**Reaction rates for every metabolic reaction in the system**

$$
\begin{aligned}
v_{R1} &= Exchange\_rate \times M8 - M1 \\
v_{R2} &= ((B + C - (B \times C)) \times M1 \times v_{max})/(M1 + K_m) \\
v_{R3} &= ((D + J - (D \times J)) \times M1 \times v_{max})/(M1 + K_m) \\
v_{R4} &= (F \times G \times M2 \times M3 \times v_{max})/(M2 + M3 + (M2 \times M3) + K_m) \\
v_{R5} &= (H \times M3 \times v_{max})/(M3 + K_m) \\
v_{R6} &= ((J + M - (J \times M)) \times M4 \times v_{max})/(M4 + K_m) \\
v_{R7} &= ((K + M - (K \times M)) \times M5 \times M4 \times v_{max})/(M5 + M4 + (M5 \times M4) + K_m) \\
v_{R8} &= ((L + M - (L \times M)) \times M6 \times M7 \times v_{max})/(M6 + M7 + (M6 \times M7) + K_m) \\
v_{R9} &= K_s \times M8
\end{aligned} \tag{S7}
$$

**Rate of change of metabolite's concentration in the system**

$$
\begin{aligned}
\frac{dM1}{dt} &= \nu_{R1} - \nu_{R2} \\
\frac{dM2}{dt} &= \nu_{R2} - \nu_{R4} \\
\frac{dM3}{dt} &= \nu_{R3} - \nu_{R4} - \nu_{R5} \\
\frac{dM4}{dt} &= \nu_{R4} - \nu_{R6} - \nu_{R7} \\
\frac{dM5}{dt} &= \nu_{R5} - \nu_{R7} \\
\frac{dM6}{dt} &= \nu_{R6} - \nu_{R8} \\
\frac{dM7}{dt} &= \nu_{R7} - \nu_{R8} \\
\frac{dM8}{dt} &= \nu_{R8} - \nu_{R9}
\end{aligned}
\tag{S8}
$$

**Rate of change of activity of the genes in the system**

$$
\begin{aligned}
\frac{dA}{dt} &= T^{-1} \times (F_{act}(M8, 1) - \frac{dA}{dt}) \\
\frac{dB}{dt} &= T^{-1} \times (F_{act}(A, 0) - B) \\
\frac{dC}{dt} &= T^{-1} \times ((F_{act}(A, 0) + F_{act}(E, 0) - (F_{act}(A, 0) \times F_{act}(E, 0))) - C) \\
\frac{dD}{dt} &= T^{-1} \times ((F_{act}(A, 0) + F_{act}(E, 0) - (F_{act}(A, 0) \times F_{act}(E, 0))) - D) \\
\frac{dE}{dt} &= T^{-1} \times (Const - E) \\
\frac{dF}{dt} &= T^{-1} \times (F_{act}(E, 0) - F) \\
\frac{dG}{dt} &= T^{-1} \times ((F_{act}(E, 0) + F_{act}(I, 0) - (F_{act}(E, 0) \times F_{act}(I, 0))) - G) \\
\frac{dH}{dt} &= T^{-1} \times (F_{act}(I, 0) - H) \\
\frac{dI}{dt} &= T^{-1} \times (Const - I) \\
\frac{dJ}{dt} &= T^{-1} \times ((F_{act}(I, 0) + F_{act}(X, 0) - (F_{act}(I, 0) \times (X, 0))) - J) \\
\frac{dK}{dt} &= T^{-1} \times ((F_{act}(I, 0) + F_{act}(X, 0) - (F_{act}(I, 0) \times (X, 0))) - K) \\
\frac{dL}{dt} &= T^{-1} \times (F_{act}(X, 0) - L) \\
\frac{dM}{dt} &= T^{-1} \times (F_{act}(X, 0) - M) \\
\frac{dX}{dt} &= T^{-1} \times (F_{act}(M8, 1) - X)
\end{aligned}
\tag{S9}
$$

### B Supplementary Figures

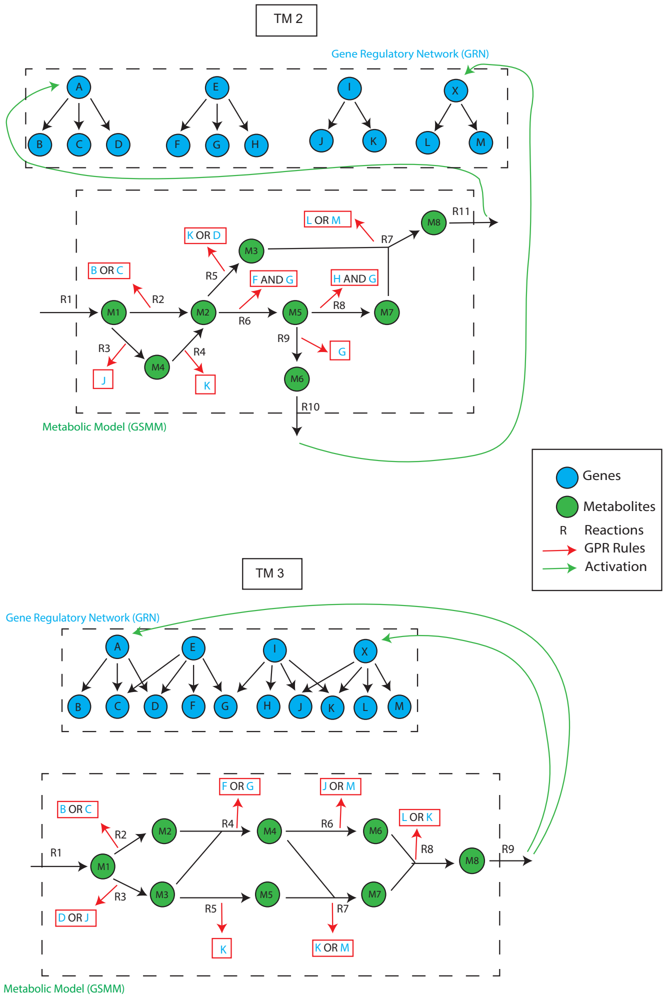

**Figure S1. Topology of TM2 and TM3:** The integrated GRN+GSMM network for TM2 and TM3 are shown.

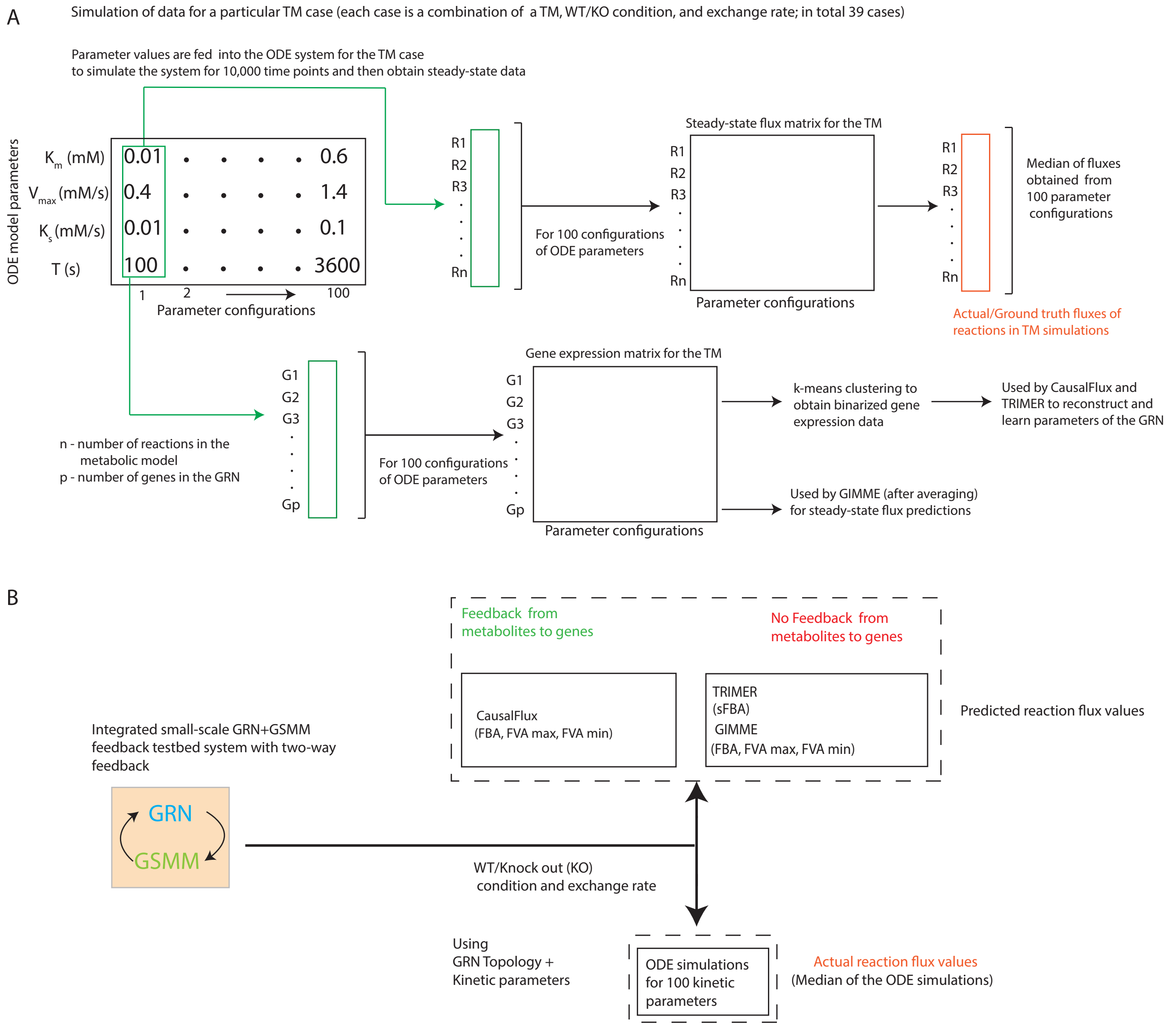

**Figure S2. Framework for simulating ground-truth fluxes/gene expression for a particular TM case; and evaluating CausalFlux, TRIMER, and GIMME on this TM case:** Consider a system of ODEs representing a given TM case, i.e., a TM (integrated small-scale GRN+GSMM testbed system with two-way feedback) under a specific WT/KO condition and a particular exchange rate. (A) For each of the 100 parameter configurations shown, we simulate this ODE system for 10,000 time points to eventually obtain steady-state data (reaction fluxes and gene expression) matrix for the given TM case. (B) Different methods (CausalFlux, TRIMER, and GIMME) are applied on the resulting steady-state data to generate predicted fluxes. The median value of steady-state fluxes across the 100 parameter configurations is taken as the actual or ground-truth fluxes. The correlation (ρ) between the predicted and ground-truth fluxes can be used to evaluate the different methods.

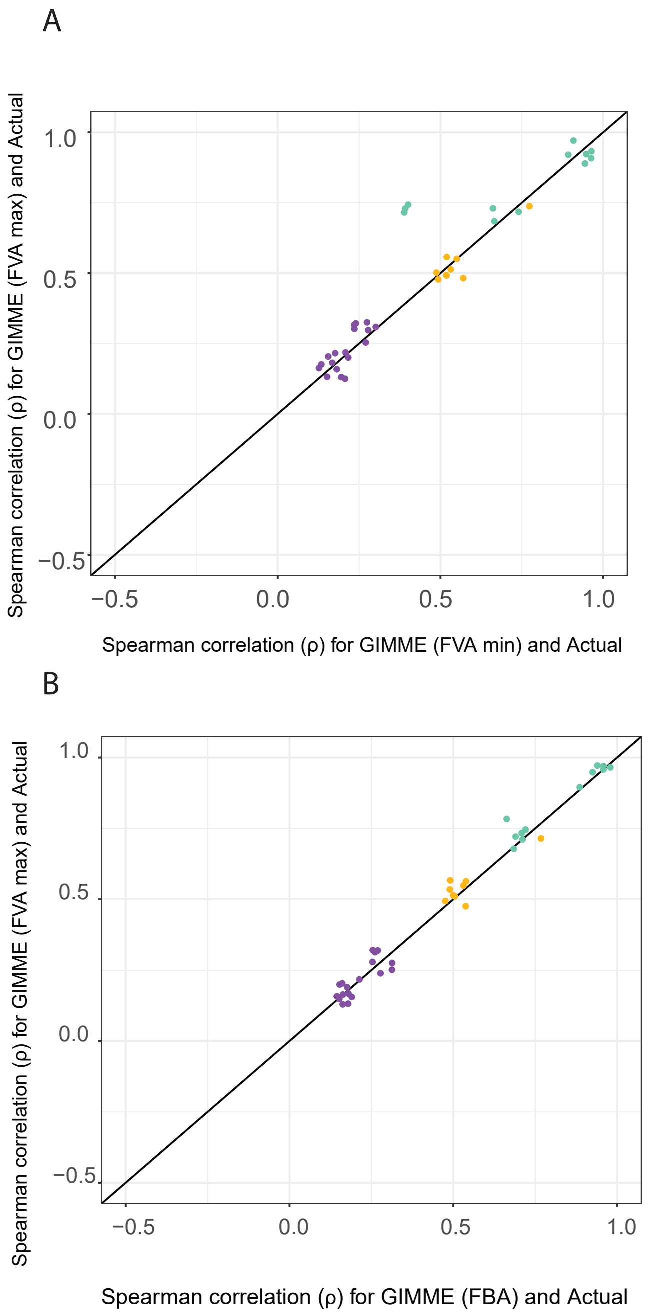

**Figure S3. Comparison of GIMME variants (FVA max, FVA min and FBA) with actual in TMs:** Scatter plot of the correlation (ρ) computed between the actual and GIMME (FVA max) for the 39 TM cases is compared with ρ computed between (A) actual and GIMME (FBA), (B) actual and GIMME (FVA min).

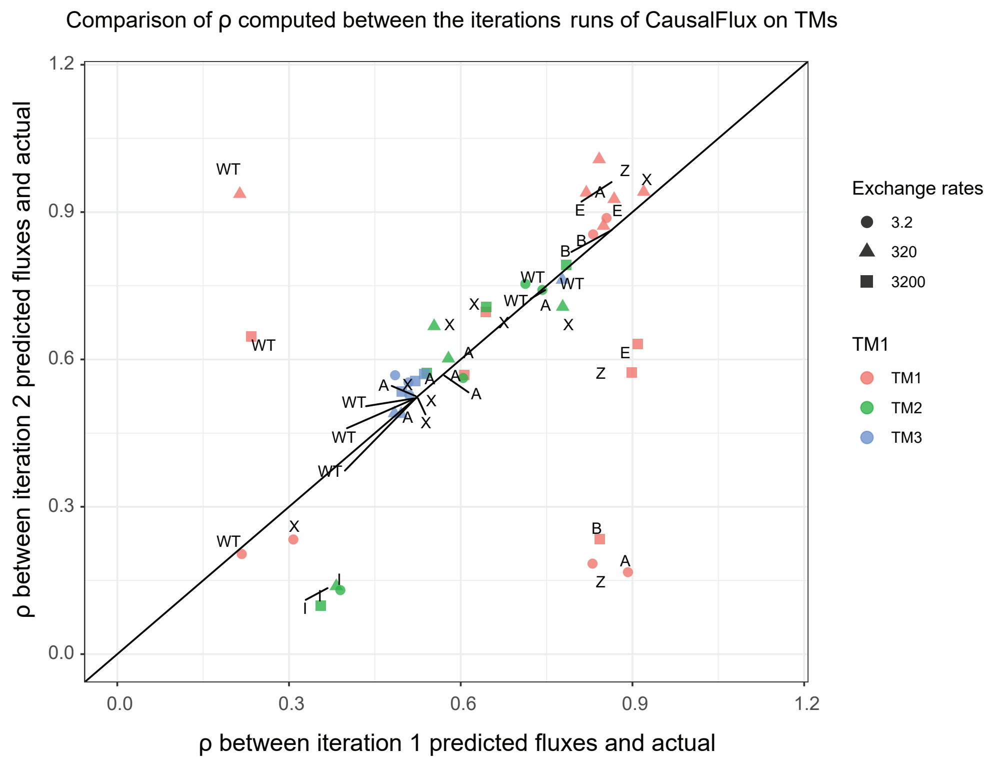

**Figure S4. Comparison of CausalFlux iteration 1 vs 2 for 39 TM cases:** Scatter plot (with jitter of 0.05 units) between the ρ computed between actual and CausalFlux predicted fluxes at iteration 1 versus iteration 2 for the 39 TM cases. The label of a point indicates its condition: "WT" for wild-type condition and the name of the gene being knocked out for single-gene KO condition.

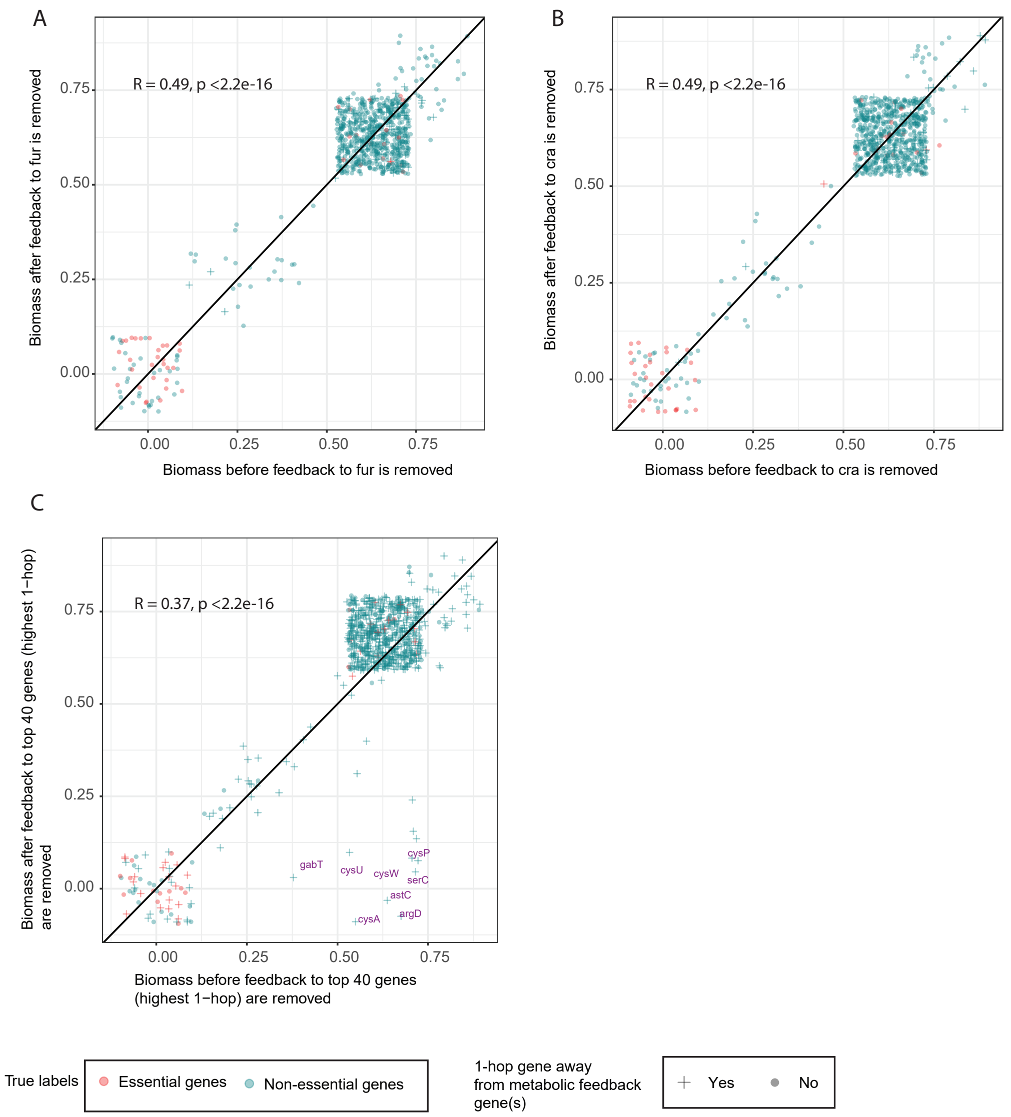

**Figure S5. Exploring the role of metabolic feedback through ablation studies:** Scatter (with default jitter) plots for ablation studies done on (A) *fur*, (B) *cra*, and (C) the set of 40 metabolic feedback genes with the most influence on single-gene KOs.

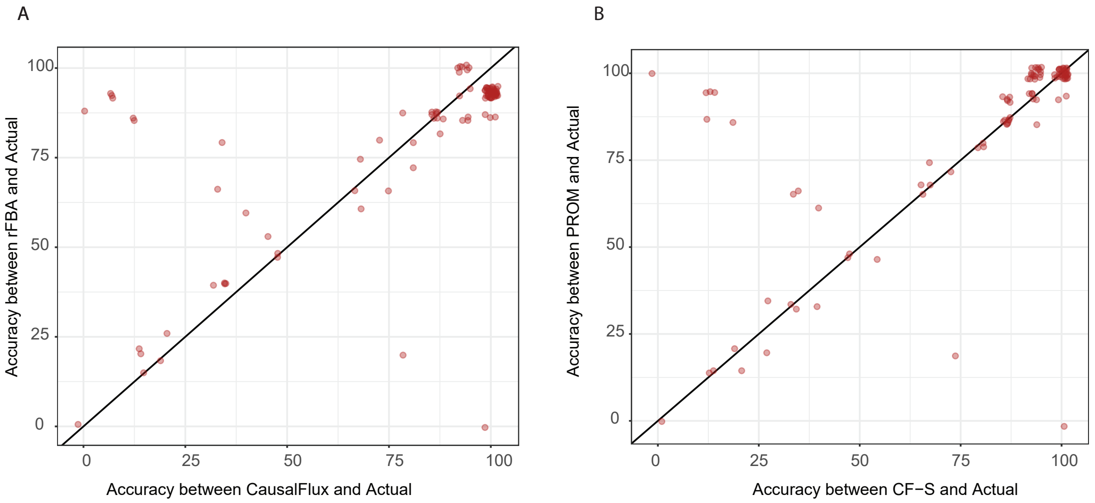

C

| Performance comparisons across media | Carbon Source (58) | Nitrogen Source (31) | Double-Nitrogen Source (11) | Overall Sources (100) |
|---|---|---|---|---|
| CausalFlux > rFBA | 43 | 8 | 5 | 56 |
| CausalFlux = rFBA | 3 | 14 | 0 | 17 |
| CausalFlux < rFBA | 12 | 9 | 6 | 27 |

D

| Performance comparisons across media | Carbon Source (58) | Nitrogen Source (31) | Double-Nitrogen Source (11) | Overall Sources (100) |
|---|---|---|---|---|
| CausalFlux > PROM | 7 | 0 | 2 | 9 |
| CausalFlux = PROM | 26 | 27 | 5 | 58 |
| CausalFlux < PROM | 25 | 4 | 4 | 33 |

E

| Methods | TP | FN | FP | TN | Precision | Recall | F1-score | Balanced Accuracy |
|---|---|---|---|---|---|---|---|---|
| CausalFlux (5 % of WT) on rFBA datasets | 150 | 214 | 105 | 1031 | 0.59 | 0.41 | 0.48 | 0.66 |
| CausalFlux (5 % of WT) on PROM datasets | 161 | 203 | 130 | 1006 | 0.44 | 0.44 | 0.49 | 0.55 |
| Random classifier | 364 | 0 | 1136 | 0 | 0.24 | 1 | 0.39 | 0.5 |
| rFBA | 258 | 106 | 187 | 949 | 0.58 | 0.71 | 0.64 | 0.77 |
| PROM | 235 | 129 | 125 | 1011 | 0.65 | 0.65 | 0.65 | 0.77 |

**Figure S6. Threshold of 5% of WT is used to define essential/non-essential cases**: (A) Scatter plot (with jitter 1.75 units) between the accuracy computed between CausalFlux predictions and Actual vs. the accuracy between rFBA predictions and Actual for the 100 media conditions. (B) Scatter plot (with jitter 1.75 units) between the accuracy computed between CausalFlux predictions and Actual vs. the accuracy between PROM predictions and Actual for the 100 media conditions. (C) Performance comparisons of CausalFlux and rFBA for the various Carbon sources, Nitrogen sources and Double-Nitrogen sources and overall 100 media conditions. (D) Performance comparisons of CausalFlux and PROM for the various Carbon sources, Nitrogen sources and Double-Nitrogen sources and overall 100 media conditions. (E) Comparison of classification metrics (TP: true positives [essential], FN: false negatives, FP: false positives, TN: true negatives [non-essential]) for CausalFlux applied to the rFBA and PROM datasets, along with Random Classifier, rFBA, and PROM predictions across 1500 data points (100 media conditions $\times$ 15 gene KOs).

A B

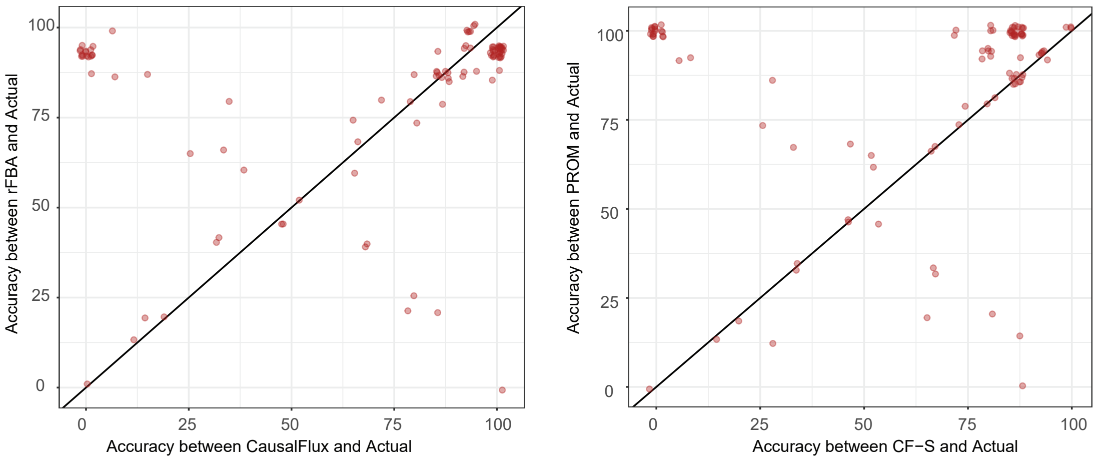

C

| Performance comparisons across media | Carbon Source (58) | Nitrogen Source (31) | Double-Nitrogen Source (11) | Overall Sources (100) |
|---|---|---|---|---|
| CausalFlux > rFBA | 31 | 8 | 5 | 44 |
| CausalFlux = rFBA | 6 | 14 | 0 | 20 |
| CausalFlux < rFBA | 20 | 9 | 6 | 36 |

D

| Performance comparisons across media | Carbon Source (58) | Nitrogen Source (31) | Double-Nitrogen Source (11) | Overall Sources (100) |
|---|---|---|---|---|
| CausalFlux > PROM | 7 | 0 | 1 | 8 |
| CausalFlux = PROM | 10 | 19 | 2 | 31 |
| CausalFlux < PROM | 41 | 12 | 8 | 61 |

E

| Methods | TP | FN | FP | TN | Precision | Recall | F1-score | Balanced Accuracy |
|---|---|---|---|---|---|---|---|---|
| CausalFlux (10 % of WT) on rFBA datasets | 265 | 99 | 398 | 738 | 0.40 | 0.73 | 0.52 | 0.69 |
| CausalFlux (10 % of WT) on PROM datasets | 280 | 84 | 502 | 634 | 0.36 | 0.77 | 0.49 | 0.66 |
| Random classifier | 364 | 0 | 1136 | 0 | 0.24 | 1 | 0.39 | 0.5 |
| rFBA | 258 | 106 | 187 | 949 | 0.58 | 0.71 | 0.64 | 0.77 |
| PROM | 235 | 129 | 125 | 1011 | 0.65 | 0.65 | 0.65 | 0.77 |

**Figure S7. Threshold of 50% of WT is used to define essential/non-essential cases**: (A) Scatter plot (with jitter 1.75 units) between the accuracy computed between CausalFlux predictions and Actual vs. the accuracy between rFBA predictions and Actual for the 100 media conditions. (B) Scatter plot (with jitter 1.75 units) between the accuracy computed between CausalFlux predictions and Actual vs. the accuracy between PROM predictions and Actual for the 100 media conditions. (C) Performance comparisons of CausalFlux and rFBA for the various Carbon sources, Nitrogen sources and Double-Nitrogen sources and overall 100 media conditions. (D) Performance comparisons of CausalFlux and PROM for the various Carbon sources, Nitrogen sources and Double-Nitrogen sources and overall 100 media conditions. (E) Comparison of classification metrics (TP: true positives [essential], FN: false negatives, FP: false positives, TN: true negatives [non-essential]) for CausalFlux applied to the rFBA and PROM datasets, along with Random Classifier, rFBA, and PROM predictions across 1500 data points (100 media conditions $\times$ 15 gene KOs).

### C Supplementary Tables

| Testbed models (TM) | Genes | Metabolites | Reactions | GPR rules | GRN (gene-to-gene edges) | metabolite-to-gene feedback edges |
|---|---|---|---|---|---|---|
| TM1 | A, B, C, D,<br>E, X, Y, Z | m1, m2, m3, m4,<br>m5 | R1: → m1<br>R2: m1 → m2<br>R3: m1 → m4<br>R4: m2 → m4 + m5 (Biomass)<br>R5: m3 → m5<br>R6: m4 →<br>R7: m5 →<br>R8: m2 → | R1: -<br>R2: C and D<br>R3: C or B<br>R4: C or D<br>R5: D or B<br>R6: -<br>R7: -<br>R8: - | Z → A<br>Y → A<br>A → C<br>B → C<br>B → D<br>E → D<br>X → B | m2 → A<br>m4 → X<br>m5 → X |
| TM2 | A, B, C, D,<br>E, F, G, H, I,<br>J, K, L, M, X | m1, m2, m3, m4,<br>m5, m6, m7, m8 | R1: → m1<br>R2: m1 → m2<br>R3: m1 → m4<br>R4: m4 → m2<br>R5: m2 → m3<br>R6: m2 → m5<br>R7: m3 + m7 → m8<br>R8: m5 → m7<br>R9: m5 → m6<br>R10: m6 →<br>R11: m8 → (Biomass) | R1:<br>R2: B or C<br>R3: J<br>R4: K<br>R5: K or D<br>R6: F and D<br>R7: L or M<br>R8: G and H<br>R9: H<br>R10:<br>R11: | A → B<br>A → C<br>A → D<br>E → F<br>E → G<br>E → H<br>I → J<br>I → K<br>X → L<br>X → M | m6 → X<br>m8 → A |
| TM3 | A, B, C, D,<br>E, F, G, H, I,<br>J, K, L, M, X | m1, m2, m3, m4,<br>m5, m6, m7, m8 | R1: → m1<br>R2: m1 → m2<br>R3: m1 → m3<br>R4: m3 → m4<br>R5: m2 → m5<br>R6: m2 → m6<br>R7: m4 + m6 → m7<br>R8: m5 + m7 → m8 (Biomass)<br>R9: m8 → | R1: -<br>R2: B or C<br>R3: D or J<br>R4: F and G<br>R5: H<br>R6: J or M<br>R7: M or K<br>R8: L or K<br>R9: - | A → B<br>A → C<br>A → D<br>E → C<br>E → D<br>E → F<br>E → G<br>I → G<br>I → H<br>I → J<br>I → K<br>X → J<br>X → K<br>X → L<br>X → M | m8 → A<br>m8 → X |

**Table S1.** For each TM, the genes, metabolites, and reactions in the TM are provided, which are in turn used to define and simulate a system of ODEs corresponding to the TM.

| **Parameters** | **Values used** |
|---|---|
| $K_m$ (mM) | 0.4, 0.5, 0.6, 0.7, 0.8, 0.9, 1.0, 1.1, 1.2, 1.3, 1.4 |
| $V_{max}$ (mM/s) | 0.01, 0.06, 0.11, 0.16, 0.21, 0.26, 0.31, 0.36, 0.41, 0.46, 0.51, 0.56 |
| $K_s$ (mM/s) | 0.01, 0.03, 0.05, 0.07, 0.09 |
| T (s) | 100, 600, 1850, 3600 |

**Table S2.** The range of parameter values used for simulating the ground truth data from the ODE equations. The parameter range shown here are for 3.2 exchange rate. $V_{max}$ and $K_s$ were scaled proportionally by 100-fold and 1000-fold for the 320 and 3200 exchange rates.

| Sl no. | Description | Number of genes | List of genes |
|---|---|---|---|
| 1 | Total predicted single-gene KOs under LB media | 63 | "nadA" "modB" "modC" "bioF" "ybjG" "dadX" "folE" "iscU" "purL" "metK" "purD" "metL" "glnA" "ilvC" "glmS" "cysG" "lysC" "ubiC" "psd" "folA" "ftsI" "mraY" "purF" "iscA" "iscS" "nadB" "fldB" "bacA" "murD" "ftsW" "dapD" "lpxD" "lpxA" "hemH" "aroA" "hemA" "ribA" "fabB" "murE" "murF" "hemL" "modA" "bioA" "bioB" "bioC" "bioD" "gltX" "purM" "murI" "glmU" "ubiA" |
| 2 | Total predicted single-gene KOs under M9 media | 133 | "dapB" "leuD" "leuB" "leuA" "murG" "murC" "ddlB" "lpxC" "nadC" "fabZ" "cynT" "aroL" "purK" "purE" "fldA" "nadA" "modB" "modC" "bioF" "ybjG" "purB" "icd" "dadX" "trpA" "trpC" "trpD" "malY" "aroH" "folE" "guaA" "iscU" "purL" "tyrA" "gabT" "cysD" "cysI" "cysJ" "argA" "lysA" "metK" "purH" "purD" "argH" "argC" "argE" "metF" "metL" "glnA" "ilvC" "ilvE" "glmS" "ilvB" "ilvN" "cysG" "lysC" "ubiC" "tyrB" "psd" "purA" "argI" "folA" "leuC" "ftsI" "mraY" "pyrC" "trpE" "purF" "cysK" "iscA" "iscS" "nadB" "cysC" "cysH" "fldB" "bacA" "ilvD" "murD" "ftsW" "dapD" "lpxD" "lpxA" "hemH" "gltA" "aroA" "hemA" "ribA" "fabB" "aroF" "metB" "pdxA" "murE" "murF" "hemL" "argF" "aroG" "modA" "bioA" "bioB" "bioC" "bioD" "trpB" "gltX" "cysM" "purC" "purM" "cysN" "argG" "murI" "argB" "glmU" "metA" "ubiA" |
| 3 | Total predicted single-gene KOs under TSB media | 93 | "dapB" "murG" "murC" "ddlB" "lpxC" "nadC" "fabZ" "cynT" "aroL" "purK" "purE" "fldA" "nadA" "modB" "modC" "bioF" "ybjG" "purB" "dadX" "folE" "guaA" "iscU" "purL" "cysD" "cysI" "cysJ" "metK" "purH" "purD" "metL" "glnA" "ilvC" "glmS" "cysG" "ubiC" "psd" "purA" "folA" "ftsI" "mraY" "pyrC" "purF" "cysK" "iscA" "iscS" "nadB" "cysC" "cysH" "fldB" "bacA" "murD" "ftsW" "dapD" "lpxD" "lpxA" "hemH" "aroA" "hemA" "ribA" "fabB" "murE" "murF" "hemL" "modA" "bioA" "bioB" "bioC" "bioD" "gltX" "cysM" "purC" "purM" "cysN" "murI" "glmU" "ubiA" |
| 4 | Common predicted single-gene KOs in all three media | 61 | "dapB" "murG" "murC" "ddlB" "lpxC" "nadC" "fabZ" "cynT" "aroL" "fldA" "nadA" "modB" "modC" "bioF" "ybjG" "dadX" "folE" "iscU" "purL" "metK" "purD" "metL" "glnA" "ilvC" "glmS" "cysG" "ubiC" "psd" "folA" "ftsI" "mraY" "purF" "iscA" "iscS" "nadB" "fldB" "bacA" "murD" "ftsW" "dapD" "lpxD" "lpxA" "hemH" "aroA" "hemA" "ribA" "fabB" "murE" "murF" "hemL" "modA" "bioA" "bioB" "bioC" "bioD" "gltX" "purM" "murI" "glmU" "ubiA" |
| 5 | Genes having growth in LB media but not in M9 media | 70 | "leuD" "leuB" "leuA" "purK" "purE" "purB" "icd" "trpA" "trpC" "trpD" "malY" "aroH" "guaA" "tyrA" "gabT" "cysD" "cysI" "cysJ" "argA" "lysA" "purH" "argH" "argC" "argE" "metF" "ilvE" "ilvB" "ilvN" "tyrB" "purA" "argI" "leuC" "pyrC" "trpE" "cysK" "cysC" "cysH" "ilvD" "gltA" "aroF" "metB" "pdxA" "argF" "aroG" "trpB" "cysM" "purC" "cysN" "argG" "argB" "metA" |
| 6 | Genes having growth in TSB media but not in M9 media | 45 | "leuD" "leuB" "leuA" "icd" "trpA" "trpC" "trpD" "malY" "aroH" "tyrA" "gabT" "argA" "lysA" "argH" "argC" "argE" "metF" "ilvE" "ilvB" "ilvN" "lysC" "tyrB" "argI" "leuC" "trpE" "ilvD" "gltA" "aroF" "metB" "pdxA" "argF" "aroG" "trpB" "argG" "argB" "metA" |

**Table S3.** List of genes that vary under different media conditions when single-gene KOs are performed on *E. coli.*

| Influence of metabolic feedback gene on the single-gene KOs | Metabolic feedback gene |
|---:|---|
| 304 | crp |
| 79 | fur |
| 56 | cra |
| 40 | lrp |
| 29 | pdhR |
| 27 | argR |
| 25 | purR |
| 21 | cysB |
| 19 | fhlA |
| 18 | nagC |
| 13 | metJ |
| 11 | paaX |
| 9 | cytR |
| 9 | argP |
| 9 | cbl |
| 9 | glpR |
| 9 | tyrR |
| 9 | malT |
| 8 | gntR |
| 8 | araC |
| 8 | trpR |
| 8 | galR |
| 8 | galS |
| 6 | allR |
| 6 | exuR |
| 6 | deoR |
| 6 | mhpR |
| 6 | lsrR |
| 5 | hcaR |
| 5 | idnR |
| 5 | uxuR |
| 5 | rbsR |
| 5 | xylR |
| 5 | fucR |
| 4 | prpR |
| 4 | caiF |
| 4 | metR |
| 4 | rhaS |
| 4 | nanR |
| 3 | rutR |
| 3 | gcvA |
| 3 | glcC |
| 3 | betI |
| 3 | cynR |
| 2 | lldR |
| 2 | iclR |
| 2 | melR |
| 2 | allS |
| 2 | treR |
| 2 | xapR |
| 1 | ilvY |
| 1 | asnC |
| 1 | nhaR |
| 0 | rhaR |

**Table S4.** For each metabolic feedback gene, this table shows its influence, i.e., the number of its 1-hop neighbors in the GRN that overlap with the set of 798 single-gene KOs in the *E. coli* dataset.

| Slno. | The gene(s) to which feedback from the metabolite(s) is removed | Precision | Recall | F1 score | Balanced Accuracy |
|---|---|---|---|---|---|
| 0 | None (Original model with all metabolite-to-gene edges intact) | 0.46 | 0.62 | 0.53 | 0.79 |
| 1 | *crp* gene | 0.41 | 0.62 | 0.49 | 0.78 |
| 2 | *fur* gene | 0.46 | 0.62 | 0.53 | 0.79 |
| 3 | *cra* gene | 0.46 | 0.62 | 0.53 | 0.79 |
| 4 | 10 metabolic feedback genes with the most influence on single-gene KOs | 0.37 | 0.62 | 0.46 | 0.77 |
| 5 | 40 metabolic feedback genes with the most influence on single-gene KOs | 0.41 | 0.62 | 0.49 | 0.78 |

**Table S5.** Performance metrics of the original model and 5 different ablation models for predicting essentiality of genes (i.e., growth vs. no-growth phenotype under single-gene KOs).

### D Supplementary Data Files

Supplementary data files listed below are available at this link: `https://github.com/BIRDSgroup/CausalFlux/tree/main/Application%20of%20CF%20on%20TMs/Applying%20methods%20on%20TMs/Supplementary%20_Data_1`

`https://github.com/BIRDSgroup/CausalFlux/blob/main/Application%20of%20CF%20on%20E.%20coli/Ablation/Supplementary_Data_2.csv`

Suppl Data 1: **Spearman's ρ between actual and predicted fluxes for all the reactions in TMs.** Spearman correlation and its p-value computed between the actual and predicted fluxes (across all the methods) for the 39 cases of TMs are given in this file.

Suppl Data 2: **Reactions with zero flux in the "gabT" and "serC" KO models with the metabolic feedback to "crp" removed.** This file lists the identifiers of the following reactions. A total of 123 reactions had zero flux, of which 27 had different GPR probabilities for the "gabT" KO model with metabolic feedback to "crp" removed. Similarly, a total of 144 reactions had zero flux, of which 29 had different GPR probabilities for the "serC" KO model with metabolic feedback to "crp" removed.
